# Intron retention as a new pre-symptomatic marker of aging and its recovery to the normal state by a traditional Japanese multi-herbal medicine

**DOI:** 10.1101/2020.02.10.941435

**Authors:** Norihiro Okada, Kenshiro Oshima, Yuki Iwasaki, Akiko Maruko, Kenya Matsumura, Erica Iioka, Trieu-Duc Vu, Naoki Fujitsuka, Akinori Nishi, Aiko Sugiyama, Mitsue Nishiyama, Atsushi Kaneko, Kazushige Mizoguchi, Masahiro Yamamoto, Susumu Nishimura

## Abstract

Intron retention (IR) is an important regulatory mechanism that affects gene expression and protein functions. Using klotho mice at the pre-symptomatic state, we discovered that retained-introns accumulated in several organs including the liver and that among these retained introns in the liver a subset was recovered to the normal state by a Japanese traditional herbal medicine. This is the first report of IR recovery by a medicine. IR-recovered genes fell into two categories: those involved in liver-specific metabolism and in splicing. Metabolome analysis of the liver showed that the klotho mice were under starvation stress. In addition, our differentially expressed gene analysis showed that liver metabolism was actually recovered by the herbal medicine at the transcriptional level. By analogy with the widespread accumulation of intron-retained pre-mRNAs induced by heat shock stress, we propose a model in which retained-introns in klotho mice were induced by an aging stress and in which this medicine-related IR recovery is indicative of the actual recovery of liver-specific metabolic function to the healthy state. Accumulation of retained-introns was also observed at the pre-symptomatic state of aging in wild-type mice and may be an excellent marker for this state in general.

## 1. Introduction

### How could we detect a pre-symptomatic marker that is associated with aging?

With demographic shifts toward older populations, it is becoming more and more important to detect pre-symptomatic markers associated with old age and to restore such a pre-symptomatic state to a healthy state so that the healthy lifespan of individuals can be prolonged by suppression of age-related diseases. It has recently become widely recognized among molecular biologists that aging reflects the gradual deterioration of the molecular components of the cell, the concerted functioning of which is vital for cell viability and proliferation (Lopez-Otin et al., 2013). In this sense, aging is regarded as a kind of disease, the extent of which varies among individuals.

The process of removing an intron from pre-mRNA is highly dynamic, during which a complex rearrangement of protein-protein, RNA-RNA, and RNA-protein interactions takes place in the spliceosome (Mayeda et al., 1994). Because of such complexity, the process is highly vulnerable to external stimuli (Mayeda et al., 1994; Padgett et al., 1986; Patel and Steitz, 2003; Stegeman and Weake, 2017) including aging, as suggested by many instances in which aging is accompanied by a change in the expression level of spliceosomal proteins or by a change in their splicing patterns (Zane et al., 2014). For example, levels of SF3B1 and SRSF family proteins can be affected by changes in alternative splicing during aging (Deschenes and Chabot, 2017). Based on these data, it is possible that some splicing-related events may function as markers for a pre-symptomatic state of aging.

### Intron retention is known to be associated with aging

Intron retention (IR) is one form of alternative splicing and has been considered to be harmful to the organism (Weischenfeldt et al., 2005) by (1) slowing down splicing kinetics to delay the onset of gene expression (Braunschweig et al., 2014), (2) increasing pre-mRNA degradation in the nucleus by nuclear exosomes (Niemela et al., 2014), and (3) increasing (pre-)mRNA degradation in the cytoplasm by nonsense-mediated decay (Lejeune and Maquat, 2005). Recent global screens of many cell and tissue types from humans and mice have, however, gradually revealed the role of IR as a negative regulator of gene expression that is integrated into a regulatory network of RNA processing and has functional significance (Jacob and Smith, 2017; Liu et al., 2017). Recent studies have shown that programmed IR is a critical regulatory pathway in normal development and differentiation, such as in granulocyte differentiation (Wong et al., 2013), terminal erythroid differentiation (Pimentel et al., 2016), male germ cell differentiation (Naro et al., 2017), and B-cell development (Ullrich and Guigo, 2019). In addition, IR accumulation has even been proposed as a post-transcriptional signature of aging (Adusumalli et al., 2019). Analyses of the transcriptome from mouse, human brain, and Drosophila head have shown a global increase in the level of IR during aging, indicating that this process may be evolutionarily conserved (Adusumalli et al., 2019). Although IR increases accumulate during aging, it is uncertain whether increasing IR of retained-introns could be a quantitative marker of the pre-symptomatic stage of aging.

### The klotho mouse is a model for premature aging

In the present study, we used the klotho mouse, which is a model for premature aging and exhibits a syndrome resembling human aging that includes a reduced lifespan, decreased activity, infertility, osteoporosis, atherosclerosis, atrophy of the skin, and emphysema (Kuro-o et al., 1997). This phenotype is caused by the disruption of a single gene, klotho (*Kl*). In a normal mouse, the klotho protein and FGF23 function together as a receptor in the kidney to exert a diuretic action on phosphorus (Razzaque, 2009). In a klotho mouse, however, the blood level of phosphorus is very high because of a deficiency in klotho protein, which leads to rapid senescence in this mouse. As aging induced by the disruption of *Kl* resembles progeria syndrome (Kuro, 2018), it does not necessarily represent normal aging. It should be noted, however, that several aspects of the *Kl* knockout mouse phenotype are known to closely represent human aging (Nabeshima, 2002; Wang and Sun, 2009).

### Juzen-taiho-to (JTT), a kind of Japanese herbal medicine, is effective in recovering physical exhaustion due to aging

Japanese multi-herbal medicines (Kampo) originated in ancient China and are widely used in Japan for various disorders. In contrast to chemically synthesized Western drugs, whose effects on humans are carefully evaluated during various stages of testing and whose mechanisms of action are generally rigorously evaluated, it has been difficult to identify the molecular substances in the Kampo that are responsible for their potency and to analyze the mechanisms by which they exert their effects. It has long been said that thousands of chemical compounds in Kampo work together, sometimes synergistically, and that their side-effects, if any, are relatively minor, although their mechanisms of action remain a total mystery (Zhou et al., 2016).

JTT, being composed of 10 crude drugs as one of the formulations approved as a medicine in Japan, was first described in a drug textbook in the Song Dynasty (AD 1151) in China and was introduced in Japan during the Kamakura period (Saiki, 2000) (AD 1185–1333). It has been traditionally administered to patients with deficiency syndrome, consisting of disturbances and imbalances in the homeostatic condition of the body (Onishi et al., 1998). More specifically, it is prescribed for patients with fatigue, anemia, night sweats, anorexia, or circulatory problems. JTT has stimulatory effects on the immune response including enhancement of phagocytosis, cytokine induction, induction of mitogenic activity of spleen cells, and activation of macrophage and natural killer cells (Matsumoto et al., 2000). JTT can also prolong life when combined with the surgical excision of tumors and has a protective effect against the deleterious effects of chemotherapy drugs (Wang et al., 2018).

### An ancient Chinese statement proposed the medicine’s usefulness for treating the pre-disease state

The ancient Chinese medical textbook *Inner Canon of the Yellow Emperor* stated that the predisease state should be treated early with Kampo medicine so that the healthy condition can be restored (UNESCO, 2011). Accordingly, ancient Chinese people knew that the healthy state can be reversibly recovered from the pre-disease state by treatment with Chinese herbal medicines. In the present study, we wanted to define this pre-disease state at the molecular level and to elucidate the efficacies of Kampo medicine by applying RNA sequencing (RNA-seq) technology and systems biology methods.

### Study hypothesis and design

We hypothesized that changes in IR patterns should be observed at the pre-symptomatic state of aging. Because it has been reported that IRs are observed during aging (Adusumalli et al., 2019), this is a reasonable assumption. Accordingly, we used klotho mice that had not yet undergone extensive aging to look for this possibility. In addition, we examined the effects of JTT on IR patterns in genes expressed in their organs by using an RNA-seq analysis (Vu et al., 2020). We discovered that IR occurred even during this earlier stage in the aging process when no changes were detected in the expression of the corresponding proteins and no prominent pathological changes in the organs were observed. Thus IR can be used as a new marker of the pre-symptomatic state.

Here we provide definitive evidence that JTT can specifically decrease IR associated with genes for liver metabolic functions among those that show an increase in IR during the process of aging. With the support of both transcriptome data related to differentially expressed genes (DEGs) and a large number of published reports about JTT efficacy with respect to liver function, we suggest that JTT has a beneficial effect on metabolism in the liver at an earlier stage of aging. We also showed that premature aging of klotho mice induces starvation-like stress. Based on these data, and based on analogy with the finding that heat shock stress induces accumulation of intron-retained mRNAs in mice (Shalgi et al., 2014), we propose a new model in which the JTT-related decrease in retained introns that accumulate in the liver at this earlier stage of aging is an indication of the effective actions of this herbal medicine at the time corresponding to the pre-symptomatic stage.

## 2. Materials and Methods

### 2.1. Animals

Male α-klotho knockout (*Kl*^−/−^/Jcl) mice (klotho mice, 3 weeks old, n = 6) and wild-type (C57BL/6JJcl) mice (WT, 3 weeks old, n = 6) were purchased from CLEA Japan, Tokyo, Japan. The mice were acclimated for 0.5 weeks in a vinyl isolator, during which they were given radiation-sterilized water and CE-2 diet (CLEA Japan) ad libitum. Thereafter, the klotho mice and WT mice were each divided into two groups, with three animals in each. In each case, one group of mice was fed CE-2 containing 0.5% (w/w) JTT, whereas the other group was fed CE-2; both groups were fed these diets from 3.5 weeks of age until the mice were 7 weeks old. As the lifespan of a klotho mouse is ~10 weeks, such mice at the end of 7 weeks are considered to be old. Accordingly, this experiment was designed to examine the effect of JTT on gene expression in the organs during the period from adolescence to senescence. Each group of mice was killed at 7 weeks of age, and their organs (blood, liver, heart and kidney) were removed and were soaked with RNA*Later* (Thermo Fisher Scientific, Rockford, IL, USA) before being subjected to RNA extraction. Tissue that was used for determining metabolites were frozen directly without soaking with RNA*Later*. Experimental procedures using animals were carried out with approval from the Laboratory Animal Committee of CLEA Japan.

### 2.2. Juzen-taiho-to (JTT)

JTT was supplied by Tsumura & Co. (Tokyo, Japan) in the form of a powdered extract. It was obtained by spray-drying a hot water extract mixture of the following 10 crude drugs in the ratios provided in parentheses: Astragali radix (10.52), Cinnamomi cortex (10.52), Rehmanniae radix (10.52), Paeoniae radix (10.52), Cnidii rhizome (10.52), Atractylodis lanceae rhizome (10.52), Angelicae radix (10.52), Ginseng radix (10.52), Poria (10.52), and Glycyrrhizae radix (5.32). The origins and species of each component, the contents of characteristic ingredients, and other pharmaceutical-grade qualities of JTT are strictly controlled as it is an ethical drug approved by the Ministry of Health, Welfare and Labor of Japan.

### 2.3. Measurement of metabolites

Approximately 50 mg of frozen tissue was added to 1,500 μL of 50% acetonitrile/Milli-Q water containing internal standards (Solution ID: 304-1002, Human Metabolome Technologies, Inc., Tsuruoka, Japan) at 0 °C to inactivate enzymes. The tissue was homogenized with three pulses at 1,500 rpm for a total of 120 sec using a tissue homogenizer (Micro Smash MS100R, Tomy Digital Biology Co., Ltd., Tokyo, Japan), and then the homogenate was centrifuged at 2,300 × *g* at 4 °C for 5 min.

Subsequently, 800 μL of the upper aqueous layer was filtered through a Millipore filter (5-kDa cutoff) at 9,100 × *g* at 4 °C for 120 min to remove proteins. The filtrate was concentrated by centrifugation and re-suspended in 50 μL of Milli-Q water for capillary electrophoresis-time-of-flight mass spectrometry (CE-TOFMS) analysis. Metabolome measurements were carried out through a facility service at Human Metabolome Technologies Inc., Tsuruoka, Japan.

### 2.4. RNA extraction and RNA-seq

RNA extraction was performed on individual tissue samples with the Pure Link RNA Mini kit (Invitrogen, MA, USA). Briefly, 600 μL of Lysis Buffer and 900 μL of TRIzol (Thermo Fisher Scientific) were added to 0.03 g of tissue, and the tissue was homogenized. After the sample was incubated for 10 min at room temperature and centrifuged at 12,000 × *g* for 15 min, the supernatant was treated with DNase and purified by column cartridge. The quality of RNA was checked with a Qubit (Thermo Fisher Scientific) and TapeStation (Agilent Technologies, CA, USA). RNA library construction was performed by using a TruSeq Stranded mRNA Sample Prep kit (Illumina, CA, USA). Paired-end (150 base pairs × 2) sequencing with the NovaSeq 6000 platform (Illumina) was outsourced to Takara Bio, Shiga, Japan.

### 2.5. Quality check and filtering of RNA-seq data and mapping analysis

For purification of the sequencing data, cutadapt v.1.16 (Deschenes and Chabot, 2017) was used to remove Illumina adapter sequences, followed by removal of the poly(A) sequence using fastx_clipper software in the fastx toolkit software package v.0.0.14 (http://hannonlab.cshl.edu/fastx_toolkit/). To remove low-quality bases or sequences, we trimmed the sequences using fastq_quality_trimmer software (parameters: -t 20 -l 30 -Q 33) and fastq_quality_filter software (parameters: -q 20 -p 80 -Q 33), both of which are included in the fastx toolkit. During the above processing, any read in which one of the pairs was missing was removed using Trimmomatic v.0.38 (Bolger et al., 2014). Then, reads containing mouse rRNA, tRNA, or phiX sequences as the control sequence from Illumina were removed using Bowtie 2 v. 2.3.4.1 (Langmead and Salzberg, 2012). We then carried out the second round of removal of any unpaired reads using bam2fastq. After completion of these filtering steps, 20 million reads of each of the forward and reverse sequences per sample were mapped to the mouse genome sequence build GRCm38 using Tophat v2.1.1 (Trapnell et al., 2009). The mouse genome sequence was downloaded from iGenomes of Illumina (http://jp.support.illumina.com/sequencing/sequencing_software/igenome.html). Multiple mapped reads were removed using samtools (parameters: samtools view -q 4). Uniquely mapped reads were counted by gene annotation (Ensembl release 81) using featureCounts v.1.6.2. The counted values were normalized with the TMM (Trimmed mean of M values) method using EdgeR library in R v.3.5.0 and used for expression analysis.

### 2.6. Analysis of alternative splicing by using rMATS

Loci with significantly different splicing patterns with *P* < 0.01 were detected within three comparisons: KL+/KL−, KL+/WT, and KL−/WT, where ‘+’ and ‘-’ refer to treatment with and without, respectively, JTT, using rMATS v.4.0.2 (Shen et al., 2014). Statistical significance was tested using the skipping junction counts (SJCs) and the inclusion junction counts (IJCs) as calculated by rMATS at the corresponding loci. We checked the mapping status and generated the mapping result view using Integrative Genomics Viewer (IGV; http://software.broadinstitute.org/software/igv/) (Robinson et al., 2011).

### 2.7. Calculation of the percent of splicing index (PSI) from mouse liver mRNA-seq data

We downloaded fastq files of mouse liver mRNA-seq data at five different timepoints (2 months, 9 months, 15 months, 24 months, and 30 months) with the ID GSE75192 from the NCBI GEO database (Aramillo Irizar et al., 2018). Based on the same procedures as those in section 2.5 described above, single-end data from 30 million reads were mapped to the mouse genome. Intron coverage and coverage of flanking exons were calculated from the BAM file of the mapping results. From genes thus characterized, we first chose 1862 genes (2787 loci) that are known to be susceptible to IR based on Ensembl transcript annotation (release 81). After removing loci with exon coverage < 5 or those with intron length < 50 base pairs, we obtained 856 loci. For the determination of the PSI, the higher value of flanking exon coverage for each gene was used to calculate its intron ratio.

### 2.8. Analysis of characteristics for the IR gene group

We analyzed enriched gene functions and pathways at 142 loci (134 genes) at which significantly different IR events were detected between KL+ and KL− using DAVID ver. 6.8 (https://david.ncifcrf.gov/) (Huang da et al., 2009) with the following criteria: Count ≥ 1, EASE ≤ 1.

### 2.9. Splice site score

For the evaluation of splice site strength, maximum entropy scores for 5’ and 3’ splice sites were calculated using MaxEntScan (Yeo and Burge, 2004).

### 2.10. Motif analysis

We compared protein-binding motifs in the mRNAs with the ATtRACT database of RNA-binding proteins and their associated motifs (Giudice et al., 2016). We compared transcription factor (TF) - binding sites using HOMER (Heinz et al., 2010) with default parameters. All detected motifs were then compared using Fisher’s test.

### 2.11. Definition of potential splicing-related genes

The KEGG pathway lists ~140 proteins as spliceosomal proteins, and we then added 109 genes from the Gene Ontology (GO) list (GO:0000398) of mRNA splicing via spliceosome. We also added 66 genes from the GO list (GO:0003729) of mRNA binding and 180 genes from the GO list (GO:0006397) of mRNA processing, both of which were chosen to overlap with genes listed by Han et al. (Han et al., 2017) that were selected based on the criterion that a knock-down experiment influenced their alternative splicing. After removing duplicates, we ultimately defined 250 genes as potential splicing-related genes in this analysis. They included almost all the spliceosomal proteins and spliceosome regulatory proteins.

### 2.12. Reverse transcription PCR amplification of IR loci

The extracted RNA was treated with Recombinant DNase I (Takara Bio, Japan) to digest the remaining genomic DNA and was purified by phenol/chloroform/isoamyl alcohol (25:24:21) and ethanol precipitation. The purified RNA was reverse transcribed using High-Capacity cDNA Reverse Transcription kits (Thermo Fisher Scientific). Primers were prepared in exons adjacent to the IR locus, and PCR amplification was performed using the cDNA. Reaction conditions were as follows: 5 μL 10× PCR buffer (Takara Bio), 5 μL dNTPs (25 mM; Takara Bio), 1 μL primers (10 pmol/μL each primer), 0.2 μL ExTaq DNA polymerase (5 U/μL; Takara Bio), 40 ng of template DNA, and DNase-free water added to a final volume of 50 μL. PCR was performed using GeneAmp PCR system 9700 (Thermo Fisher Scientific) with the following conditions: initial annealing at 96 °C for 5 min, followed by 25 or 30 cycles (96 °C for 30 sec, 55 °C for 45 sec, 72 °C for 2 min). After a final extension at 72 °C for 5 min, the PCR mixtures were held at 4 °C. The amplicon was confirmed based on size by TapeStation. Primer sequences are listed in Table S1.

### 2.13. Analysis of DEGs

Genes that had significantly differential expression with *P* < 0.01 in two comparisons, namely KL+/KL− and KL−/WT, were detected using edgeR (Robinson et al., 2010). The recovered genes that were significantly down-regulated in KL− as compared with WT and significantly up-regulated in KL+ as compared with KL– were extracted.

### 2.14. Western blot analysis

Mouse liver samples were homogenized in ice-cold RIPA buffer (50 mM Tris, 150 mM NaCl, 1% NP-40, 0.5% deoxycholic acid sodium monohydrate, 0.1% SDS, 10 mM NaF, pH 7.4) with a disposable homogenizer (Nippi BioMasher, Nippi, Inc. Tokyo, Japan) (30 strokes), and the homogenates were centrifuged at 10,000 × *g* and 4 °C for 20 min. The protein concentration of each supernatant was measured by the BCA protein assay (Thermo Fisher Scientific), and supernatants were diluted to equal protein concentrations, combined with 2 M DTT (final concentration, 0.2 M) and 4× SDS sample buffer (6% SDS, 40% glycerol, 0.4% bromophenol blue, 250 mM Tris, pH 6.8), and boiled for 5 min at 95 °C.

Protein samples (30 μg/well) were resolved by SDS-PAGE and then transferred onto PVDF membranes. The membranes were blocked with 5% milk in TBST (20 mM Tris, 150 mM NaCl, containing 0.05% Tween-20, pH 7.4) and incubated with primary antibodies at 4 °C overnight. Primary antibodies used in this study were anti-RGN/SMP30 (17947-1-AP, Proteintech), anti-DDX5, p68 (10804-1-AP, Proteintech), anti-NDUFS2 (GTX114924, GeneTex), anti-PPARD (10156-2-AP, Proteintech), anti-SIRT7 (12994-1-AP, Proteintech), anti-NXF1 (10328-1-AP, Proteintech), anti-SRSF6 (11772-1-AP, Proteintech), anti-LXR beta (ab28479, Abcam), and anti-GAPDH (sc-32233, Santa Cruz Biotechnology). After incubation with secondary antibodies peroxidase-conjugated anti-rabbit IgG (SA00001-2, Proteintech) or anti–mouse IgG (sc-516102, Santa Cruz Biotechnology), protein bands were detected using ECL Prime Western Blotting Detection Reagents (GE Healthcare). Some of the membranes were probed with anti-GAPDH, which was used as the loading control for other blots in each experiment. The signal intensity was quantified using ImageJ (NIH) or a ChemiDoc system (Bio-Rad). Western blots were repeated a minimum of three times with different animals, and representative blots are shown.

## 3. Results and Discussion

### 3.1. Identifying a marker of the pre-symptomatic state in klotho mice at 7 weeks of age

We used klotho mice, a model for premature aging similar to progeria in humans, to look for a new marker for the pre-symptomatic state of aging as well as the possible effect of a Japanese herbal medicine on this state. Because aging is currently regarded as a kind of disease (see Introduction), a related pre-symptomatic state present before notable aging occurs is critical for medical treatment. We chose the timepoint of 7 weeks after birth for the date of sampling (Figure 1A), which is less than half of the half-life of these mice (14.7 weeks; Figure 1B). We expected that any markers isolated from these mice at 7 weeks would represent the pre-symptomatic state. We also designed our experiments to examine the effect of JTT on the improvement of this state (Figure 1A). The klotho mice (KL) and wild-type mice (WT) were fed with (+) or without (-) 0.5% JTT from 3.5 weeks of age until the mice were 7 weeks old. Consistent with our knowledge that Kampo medicine does not have a biological effect on a healthy recipient (Goswami et al., 2019; Shimada et al., 2008), the data for WT fed with and without JTT were essentially the same. Accordingly, the data for WT mice fed with JTT are not shown in the main text, although subsets are shown in the supplementary figures. By 7 weeks of age, the mouse liver is regarded as being in the adult stage in terms of its transcriptome (Gunewardena et al., 2015; Kadota et al., 2020), justifying the use of WT mice at this age as the control.

**Figure 1.**
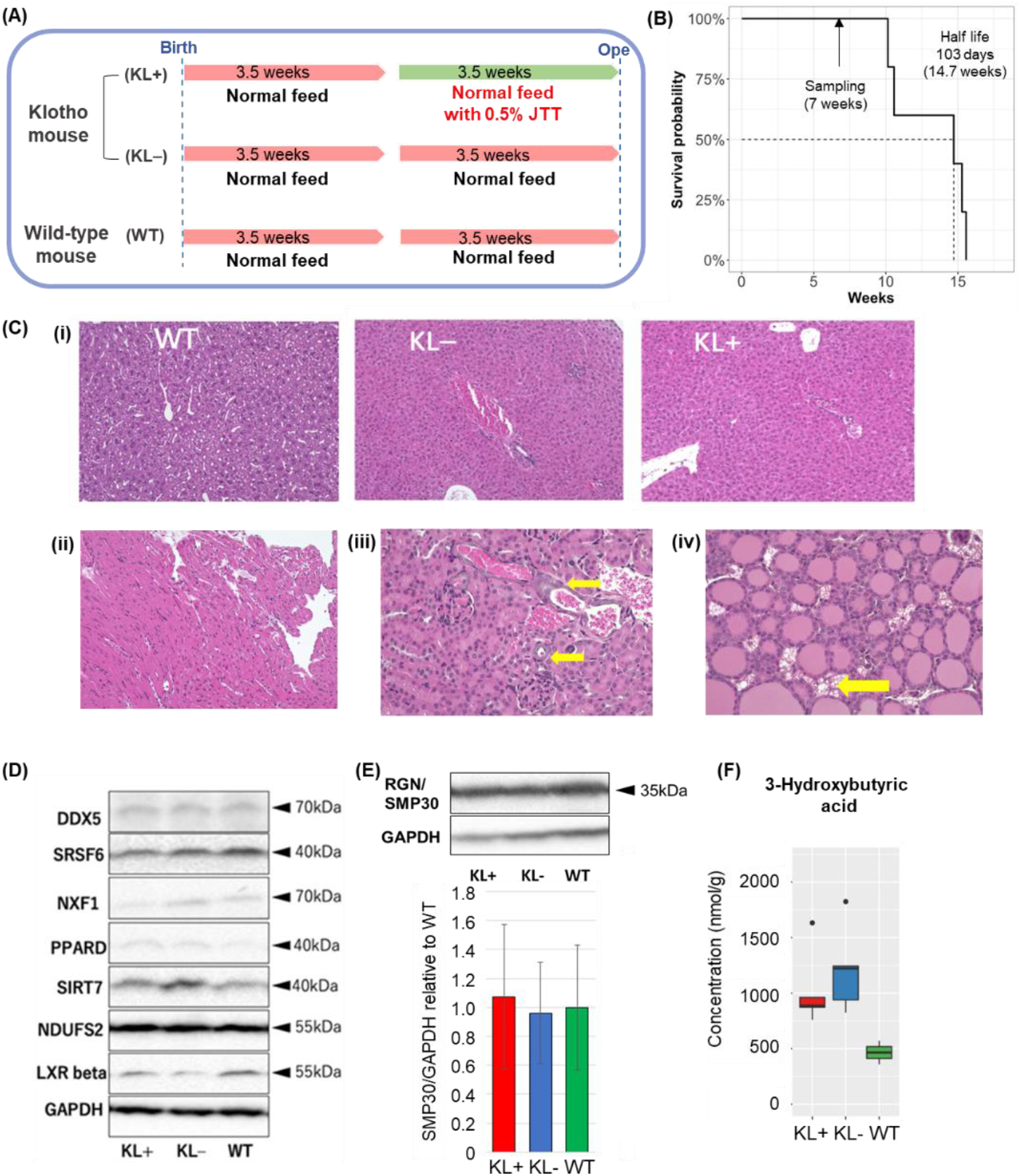
The pre-symptomatic state of klotho mice at 7 weeks of age. (A) Schematic illustration of animal experiments. Three-week-old male α-klotho knockout (*Kl^−/−^*/Jcl) mice (klotho mice, n = 8) and WT (C57BL/6JJcl) mice (n = 8) were purchased from CLEA Japan, Tokyo, Japan. The mice were acclimated for 0.5 weeks in a vinyl isolator, during which they were given radiation-sterilized water and CE-2 diet (CLEA Japan) ad libitum. Thereafter, the klotho mice were divided into two groups, with four animals in each. In each case, one group of mice was fed CE-2 containing 0.5% (w/w) JTT, whereas the other group was fed CE-2 only from 3.5 weeks of age until the mice were 7 weeks old. ‘Ope’ stands for operation. (B) Estimation of the half-life of klotho mice (n = 5) by survival-curve analysis. (C) Histological observations of (i) liver tissue from 7-week-old KL+ and KL− showed that it did not exhibit any signs of aging relative to WT mice, as was also true for (ii) heart tissue. Histological observations of tissue from (iii) kidney and (iv) thyroid showed that these organs had begun to exhibit a senescent phenotype at this age. Yellow arrows indicate such phenotypic changes (see text). (D) The expression of seven proteins, each of which showed IR, was examined by western blotting using liver tissues. Expression levels did not change in KL+, KL−, and WT at 7 weeks of age. (E) Representative western blotting data (upper) of a senescence marker protein, RGN/SMP30, together with its quantified expression (lower) show that KL− are not senescent. Statistical significance was estimated by one-way ANOVA. Error bars indicate the mean ± standard deviation of triplicate measurements. (F) An analysis for 3-HBA showed that the KL− are undergoing a starvation-like condition at this age.

First, to examine the extent of aging in liver tissue from klotho mice, we looked for histological changes at 7 weeks of age relative to liver tissue from WT mice. Figure 1C(i) shows that there were no alterations in the liver tissue, such as calcification, indicating that the liver did not exhibit a senescent phenotype at this age. We also examined tissue from the heart (Figure 1C(ii)), kidney (Figure 1C(iii)), and thyroid (Figure 1C(iv)) as controls. Heart tissue from KL mice looked normal, but tissue from the kidney and thyroid showed evidence of senescence based on calcification and accumulation of vacuoles among calcitonin-producing cells, respectively.

Second, we chose several proteins and examined their levels by performing western blotting analysis using liver tissue (Figure 1D & S1). The representative proteins are DDX5, SRSF6, and NXF1, all of which are splicing-related genes, and PPARD and SIRT7, both of which are involved in metabolic pathways of the liver. NDUFS2 is the core subunit of the mitochondrial Complex-1, and LXRβ is encoded by *Nr1h2* and regulates macrophage function. GAPDH was used as the loading control. Furthermore, we checked the expression of RGN/SMP30, a marker of senescence that shows decreased expression with aging (Maruyama et al., 2010) (Figure 1E). There was no observable difference in the expression of these proteins between KL− and WT (Figure 1D & 1E), and thus we can conclude that the klotho mouse liver at 7 weeks of age does not show clear signs of aging.

### 3.2. Metabolome analysis shows that klotho mice at 7 weeks of age express starvation-like characteristics

To look for possible internal changes in the liver that could describe the pre-symptomatic state in this earlier stage of aging, we investigated 110 metabolites of the liver using CE-TOFMS and compared metabolite levels between KL− and WT, the comparison of which is a measure of premature aging. Among 12 metabolites that showed such differences (*P* < 0.05; Figure S2), it is remarkable that 3-hydroxybutyric acid (3-HBA) was significantly increased in KL− (Figure 1F), to a level 2.5 times higher than that in WT. 3-HBA is synthesized in the liver from fatty acid as one of the ketone bodies under conditions of starvation and can be used as an energy source when blood glucose is low (Fukao et al., 2014). Consistent with this, certain intermediate metabolites of glycolysis, namely glucose 1-phosphate, fructose 1,6-diphosphate, dihydroxyacetone phosphate, lactic acid, CoA divalent, and alanine in KL− were significantly decreased as compared with their levels in WT (Figure S2). These data suggest that klotho mice are undergoing a starvation-like state, in spite of the presence of ample food, during which the synthesis of 3-HBA is activated. Accordingly, an increase in 3-HBA and a reduction in glycolysis in KL− might be representative of premature aging in klotho mice.

### 3.3. DEG analysis shows the alteration of gene expression during premature aging

Knowing that the liver of klotho mice is under this starvation-like condition, we performed RNA-seq analysis to look for a possible alteration of the gene expression level in the liver during the aging process. From liver collected from KL+, KL−, and WT, we isolated mRNAs and then sequenced them. Multi-dimensional scaling confirmed the consistency of each expression dataset (Figure S3A). In addition, transcriptomes from individual organs from KL+ and KL− indicated that JTT induced extensive transcriptome changes (Figure S3B). We then extracted the set of down-regulated genes and the set of up-regulated genes in KL− from the comparison of KL− and WT (1228 genes and 1352 genes, respectively; Figure 2A). The former represents genes whose expression decreased in the liver probably due to the premature aging-related deterioration, and the latter represents genes whose expression increased, possibly reflecting an adaptive response to this aging. To characterize enriched cell types in these genes, we performed a cell type enrichment (Cten) analysis (Shoemaker et al., 2012) by using the software provided by these authors, in which, for each cell type, a transcript was defined as a highly expressed cell-specific gene in that cell if it had an expression value at least 15 times greater than its median expression value across all cells examined. The software can accurately identify the appropriate cell type in a highly heterogeneous environment and provides insight into the suggested level of enrichment to minimize the number of false discoveries. Figure 2C shows that liver-specific genes were especially enriched among the genes whose expression decreased during the premature aging, suggesting that liver function was reduced during aging. In contrast, a variety of non-liver-specific cell types were enriched among those genes whose expression increased during the premature aging (Figure 2D). It should be noted that several macrophage-related cells were enriched in these increased genes, consistent with the generally believed notion that the aging process activates innate immunity in spite of its severe deleterious effects on adaptive immunity (Solana et al., 2006).

**Figure 2.**
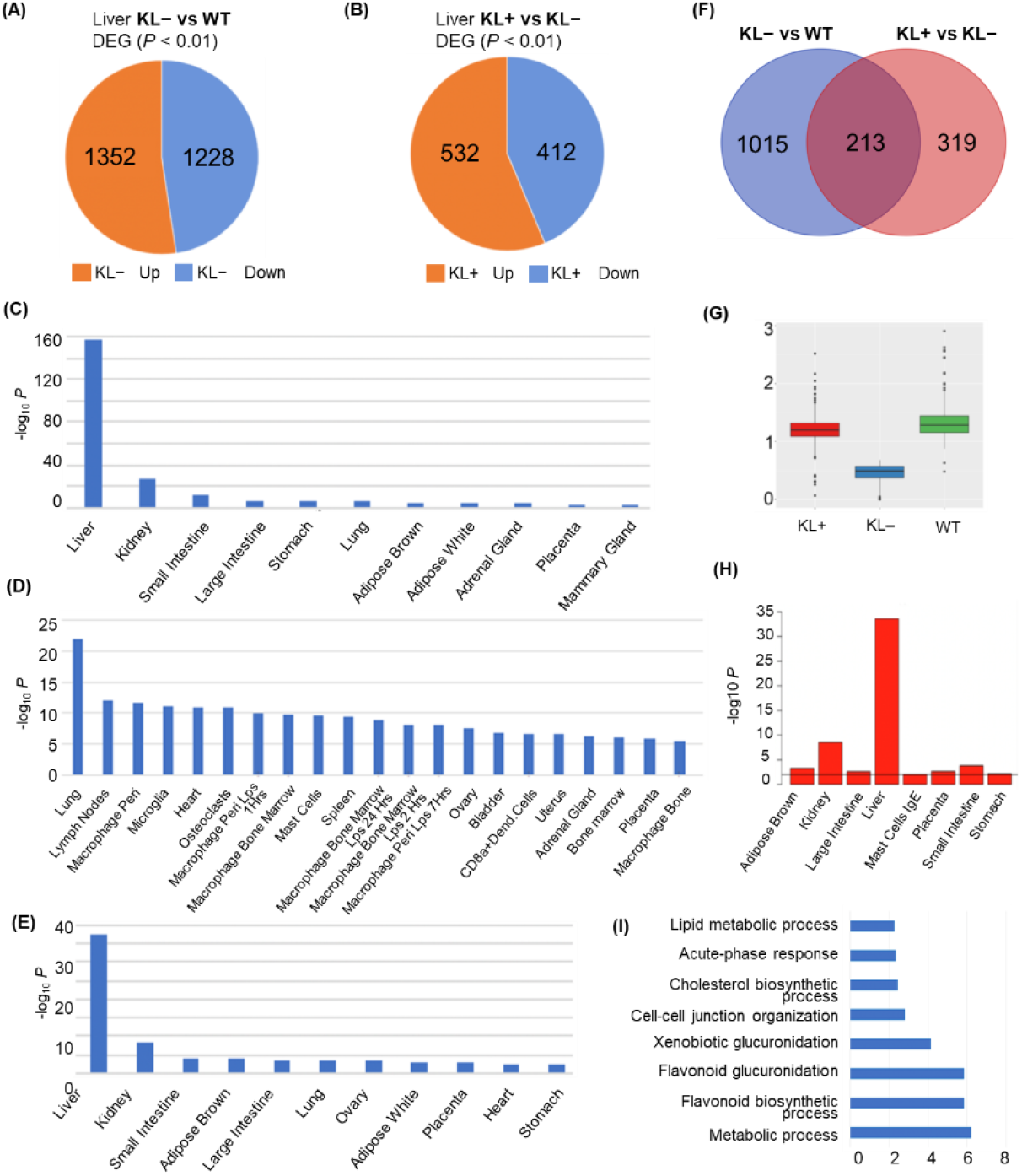
DEG analysis showed that liver function in klotho mice improved after administration of JTT. Number of up-regulated genes and down-regulated genes from the DEG data for the comparison (A) of KL− vs WT and (B) of KL+ vs KL−. Cten analysis (http://www.influenza-x.org/~jshoemaker/cten/) of (C) 1228 down-regulated genes in KL− in the comparison of KL− vs WT, (D) 1352 up-regulated genes in KL− in the comparison of KL− vs WT, and (E) 532 up-regulated genes in KL+ in the comparison of KL+ vs KL−. (F) A Venn diagram of transcriptome comparisons from KL+ vs. KL− and KL− vs. WT. Two hundred thirteen genes were shared between the comparisons. (G) Boxplot of expression data for the 213 genes shows a significant decrease in KL− in comparison with that in WT and a significant increase in KL+ in comparison with KL−. Values along the vertical axis are normalized based on the overall average. (H) Cten analysis for the 213 genes that were differentially expressed in the comparison between KL+ vs KL− and between KL− vs WT. The horizontal line indicates *P* = 0.01. (I) Bar graph showing −log_10_ *P* of GO terms enriched for the 213 genes shown in (F). Only those terms with *P* < 0.01 are shown.

### 3.4. DEG analysis shows an improvement in liver function as a result of JTT administration

To examine the influence of administration of JTT on the gene expression level in the livers of klotho mice, we extracted the set of genes that were up-regulated and down-regulated in KL+ after administration of JTT from the comparison of KL+ and KL– (532 genes and 412 genes, respectively; Figure 2B). Cten analysis of the up-regulated genes demonstrated that liver-specific genes were highly enriched (Figure 2E), whereas the down-regulated genes did not show any such enrichment (data not shown). The up-regulated gene set in KL+ from the comparison of KL+ and KL– (532 genes) and the down-regulated gene set in KL– from the comparison of KL– and WT (1228 genes), both of which showed liver-specific enrichment by Cten analysis, were compared, resulting in 213 commonly shared DEGs, all of which exhibited a clear recovery pattern from the decreased level of KL– to the level of WT in KL+ (Figure 2F). Figure 2G shows a boxplot summarizing the expression of 213 genes, in which each data point was normalized to the average expression level of each gene, and all the data were summed. We used these genes for a Cten analysis and showed that liver-specific functions were highly enriched in the 213 genes (Figure 2H). We then extracted the GO terms that were enriched for these genes. Figure 2I shows that genes related to liver-specific functions, including metabolic process, flavonoid biosynthetic process, flavonoid glucuronidation, and lipid metabolic process, were enriched. As liver-specific genes related to the metabolism of glucose, lipids, amino acids, and cholesterol are controlled at the level of transcription (Desvergne et al., 2006; Yabaluri and Bashyam, 2010), it can be said that liver function, which had been weakened by down-regulation of liver-specific metabolic genes associated with aging, was recovered at the level of transcription by administration of JTT. The many reports showing that JTT is effective for the recovery of liver damage (see section 3.10) are also consistent with these data.

### 3.5. IR and its recovery after administration of JTT are most prominent in the liver

Knowing that klotho mice at 7 weeks of age represent a starvation-like state and that their liver-specific functions at that time might be reduced due to the decreased expression of liver-specific genes, we looked for possible changes in alternative splicing in the liver at this age because it is known that several stresses can change alternative splicing patterns (Boutz et al., 2015; Ninomiya et al., 2011; Shalgi et al., 2014) (see the discussion below). We performed RNA-seq analysis in five additional organs, namely, blood, bone, brain, heart, and kidney, for comparison.

To identify what type of alternative splicing is most prominent in the six organs examined here, we first analyzed five types of alternative splicing—alternative 5’ splice site, alternative 3’ splice site, skipped exon (SE), IR, and mutually exclusive exons—between KL+ and KL– by using rMATS (Table 1-1). SE and IR were the most prominent types of alternative splicing among the six organs. Noticeably, changes in the frequency of IR in the liver and bone were most notable, followed by the change in the IR frequency in the blood (Table 1-1). We then calculated the difference in the frequency of IR in the liver in comparisons among KL+, KL–, and WT with the criteria of *P* < 0.01 (Table 1-2). A difference in the frequency of retained introns for the comparison between KL+ and WT was noted for only 59 loci, whereas that for KL– and WT was noted for 212 loci. The comparison between KL+ and WT is a measure of the effect of JTT on senescence, whereas the comparison between KL– and WT measures the effect of the senescence level in terms of the extent of retained introns accumulated during premature aging in klotho mice. The above data suggest that the administration of JTT shifted the alternative splicing pattern in klotho mice to be more similar to that of WT mice.

**Table 1-1.**
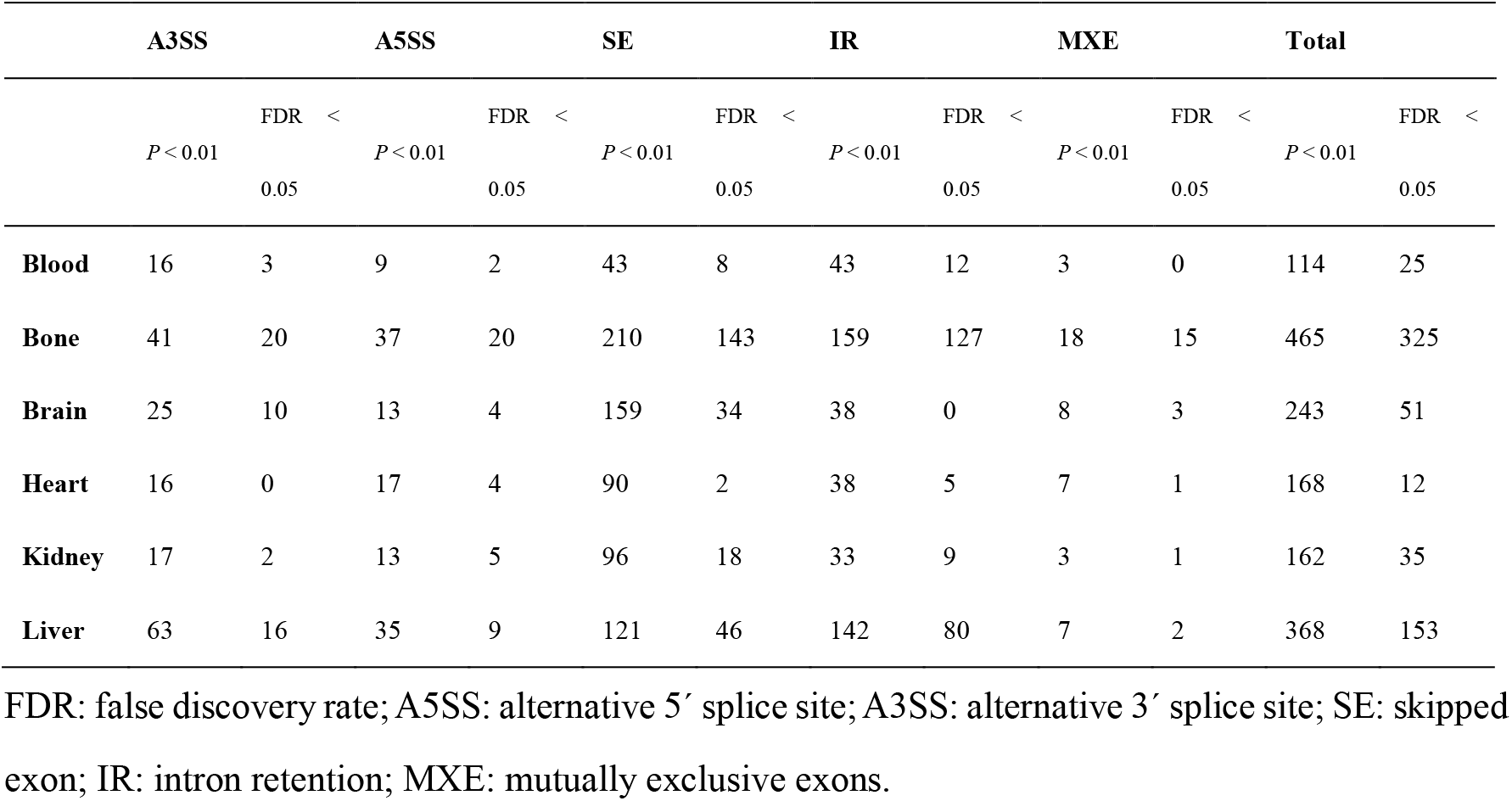
Number of alternative splicing loci with significant differences between KL+ and KL–.

|  | A3SS |  | A5SS |  | SE |  | IR |  | MXE |  | Total |  |
| --- | --- | --- | --- | --- | --- | --- | --- | --- | --- | --- | --- | --- |
| | $P < 0.01$ | FDR < 0.05 | $P < 0.01$ | FDR < 0.05 | $P < 0.01$ | FDR < 0.05 | $P < 0.01$ | FDR < 0.05 | $P < 0.01$ | FDR < 0.05 | $P < 0.01$ | FDR < 0.05 |
| **Blood** | 16 | 3 | 9 | 2 | 43 | 8 | 43 | 12 | 3 | 0 | 114 | 25 |
| **Bone** | 41 | 20 | 37 | 20 | 210 | 143 | 159 | 127 | 18 | 15 | 465 | 325 |
| **Brain** | 25 | 10 | 13 | 4 | 159 | 34 | 38 | 0 | 8 | 3 | 243 | 51 |
| **Heart** | 16 | 0 | 17 | 4 | 90 | 2 | 38 | 5 | 7 | 1 | 168 | 12 |
| **Kidney** | 17 | 2 | 13 | 5 | 96 | 18 | 33 | 9 | 3 | 1 | 162 | 35 |
| **Liver** | 63 | 16 | 35 | 9 | 121 | 46 | 142 | 80 | 7 | 2 | 368 | 153 |

FDR: false discovery rate; A5SS: alternative 5′ splice site; A3SS: alternative 3′ splice site; SE: skipped exon; IR: intron retention; MXE: mutually exclusive exons.

**Table 1-2.** Number of alternative splicing loci with significant differences in the liver.

|  | SE | A5SS | A3SS | MXE | IR | Total |
| --- | --- | --- | --- | --- | --- | --- |
| **KL+/KL-** | 121 | 35 | 63 | 7 | 142 | 368 |
| **KL+/WT** | 199 | 31 | 43 | 7 | 59 | 339 |
| **KL-/WT** | 205 | 36 | 77 | 9 | 212 | 539 |

SE; skipping exons, A5SS; alternative 5' splice site, A3SS; alternative 3' splice site, MXE; mutually exclusive exons, IR; intron retention

The 142 IR loci (Table 1-1 & 1-2) that were differentially affected between KL+ and KL– in the liver included those genes for which the frequency of IR was decreased with JTT in KL+ (132 loci; designated as “DecIR” type) and those genes for which the frequency of IR was increased with JTT in KL+ (10 loci; designated as “IncIR” type) (Figure 3A). Likewise, the comparison between KL+ and KL– for the other five organs showed a number of differential IR loci that could be divided into these two types. The liver and bone as well as blood exhibited higher values in the DecIR type as compared with the IncIR type, whereas the other three organs, brain, kidney, and heart, had comparable values for DecIR and IncIR (Figure 3A). From these data, we designated liver and bone as well as blood as three target organs for JTT. As we showed that liver-specific function was recovered by administration of JTT (see section 3.4), we focused in particular on the analyses of genes that underwent a decrease in IR in the liver in response to JTT.

**Figure 3.**
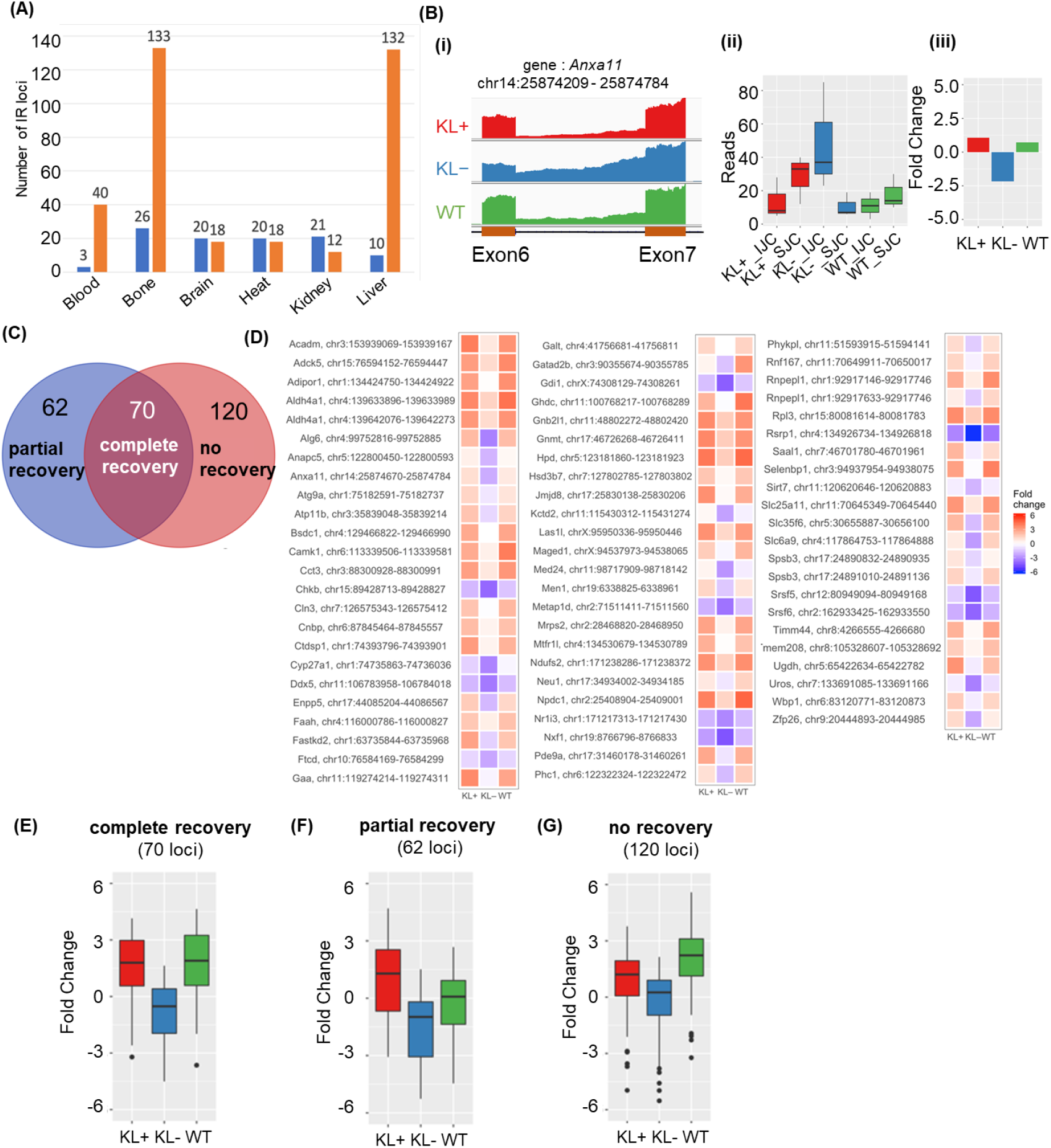
Quantification of IR events. (A) Among the six organs examined, liver and bone had the most notable recovery of retained introns after administration of JTT. Bar graphs show the number of retained introns that decreased (in orange) or increased (in blue) in the organs as a result of administration of JTT. Based on the rMATS analysis, genes with a significant change in IR in the comparison of KL+ vs KL– were divided into two groups. In the liver, the first group included genes that showed a decrease in IR in KL+ (132 loci; DecIR type, implying that the FC of KL+ was larger than that of KL–), and the second included genes with an increase in IR in KL+ (10 loci; IncIR type, implying that the fold change of KL+ was smaller than that of KL–). Similarly, in the bone, the first group included genes that showed a decrease in IR in KL+ (133 loci; DecIR type), and the second included genes with an increase in IR in KL+ (26 loci; IncIR type). (B) (i) The mapping results from KL+, KL−, and WT are shown for a single gene, *Anxa11*, using IGV Thick black bars indicate the sites of junctions in the reads where SJCs and IJCs were analyzed. (ii) Read counts from KL+, KL−, and WT for IJCs and SJCs. (iii) FC in the number of SJCs relative to IJCs for KL+, KL−, and WT. (C) A Venn diagram of two types of loci, the DecIR type (blue circle) in the comparison of KL+ and KL− and the IncIR type (red circle) in the comparison of KL− and WT. Three categories of loci, namely “partial recovery”, “complete recovery”, and “no recovery”, are distinguishable. (D) Heatmap illustration of FC values of loci in KL+, KL−, and WT for the 70 “complete recovery” loci. (E-G) Boxplots of the FC values of (E) the 70 “complete recovery” loci, (F) the 62 “partial recovery” loci, and (G) the 120 “no recovery” loci in KL+, KL−, and WT.

### 3.6. Analysis of IR-recovered genes in the liver

To demonstrate our strategy for evaluating the extent of IR, we first show an example based on the data for *Anxa11* expression in the liver (Figure 3B). *Anxa11* plays an important role in cell division, Ca^2+^ signaling, vesicle trafficking, and apoptosis (Han et al., 2017). Sequenced *Anxa11* transcripts showed extensive accumulation of a retained intron in KL−, whereas this intron was retained to a lesser extent for KL+ and WT (Figure 3B(i)).

Then, to analyze IR of *Anxa11* mathematically, we calculated the read counts of skipping junctions (SJCs) and those of inclusion junctions (IJCs) for KL+, KL−, and WT (Figure 3B(ii)) according to the strategy illustrated by rMATS (Figure S4A). For KL+, we then took the base-2 logarithm of the SJC/IJC ratio, which gives the fold change (hereafter designated as FC) between the SJC and IJC at the intron junction (the thick black bars in Figure 3B(i)). The same process was carried out for KL− and WT, and the FC values for KL+, KL−, and WT were compared. The lower FC value for KL− was recovered in KL+ to the same level as that of WT (Figure 3B(iii)). The V shape of this bar plot indicated a recovery from the senescence-type IR to the healthy-type IR after JTT administration (see below).

As was described above, we first obtained the DecIR type (132 loci, 122 genes) in klotho mice caused by JTT administration by comparing KL+ vs KL−. This simply represents the recovery of retained introns in KL+. Then, we obtained the IncIR type (190 loci, 179 genes) in klotho mice by comparing KL− vs WT. This represents a measure of the progeria-like senescence-related changes in retained introns. Accordingly, the comparison between these two types allowed us to highlight the effect of JTT at 7 weeks of age as shown in a Venn diagram (Figure 3C). The overlapping 70 loci (67 genes) represent the “complete recovery” loci, which are defined as those with significant differences between KL+ and KL− and between KL− and WT, but with FC values for KL+ and WT that were similar. The non-overlapping 62 loci (61 genes) and 120 loci (115 genes) represent “partial recovery” and “no recovery” loci, respectively. Figure 3D shows a heatmap of the 70 “complete recovery” loci, each of which represents a clear recovery pattern. Figure 3E shows a boxplot of the FC values of KL+, KL−, and WT for these 70 loci, which shows the typical V-shaped pattern of recovery. Figure 3F & 3G show boxplots of the “partial recovery” loci and the “no recovery” loci, respectively. The former represents loci with a significant difference between KL+ and KL− and without a significant difference between KL− and WT. The latter represents loci with a significant difference between KL− and WT and without a significant difference between KL+ and KL−.

### 3.7. Characteristics of loci with retained introns in the liver

We next characterized introns among the 70 “complete recovery” loci, the 120 “no recovery” loci and all other genes (~250,000 loci) expressed in the liver. The average length of introns from the “complete recovery” and “no recovery” loci was significantly shorter than that of all other introns (Figure 4A). There was, however, no significant difference in intron length between the “complete recovery” and “no recovery” loci, suggesting that the “complete recovery” loci were not selectively discriminated from the “no recovery” loci during the recovery process associated with JTT administration. The GC content of introns associated with the “complete recovery” and “no recovery” groups was significantly higher than that of all other liver-related introns (Figure 4B). Also, there was no significant difference between “complete recovery” and “no recovery” introns in this respect, suggesting that GC content was not linked to recovery status among IR loci. Then, we examined the strength of 5’ and 3’ splice sites by calculating scores for these splice sites using the software MaxEntScan. The 5’ splice sites of introns of the “complete recovery” and “no recovery” loci were slightly but significantly weaker than those of all other liver-related introns (Figure 4C), but we found no such differences at the 3’ splice sites (Figure 4D). These characteristics of introns at IR loci are consistent with previous data (Braunschweig et al., 2014). We also confirmed the susceptibility of introns to IR in genes in terms of their shorter intron length and higher GC content in the other two organs, and weaker intron score at the 5’ and 3’ splice sites in the brain (Figure S5) as was shown in the case of liver (Figure 4).

**Figure 4.**
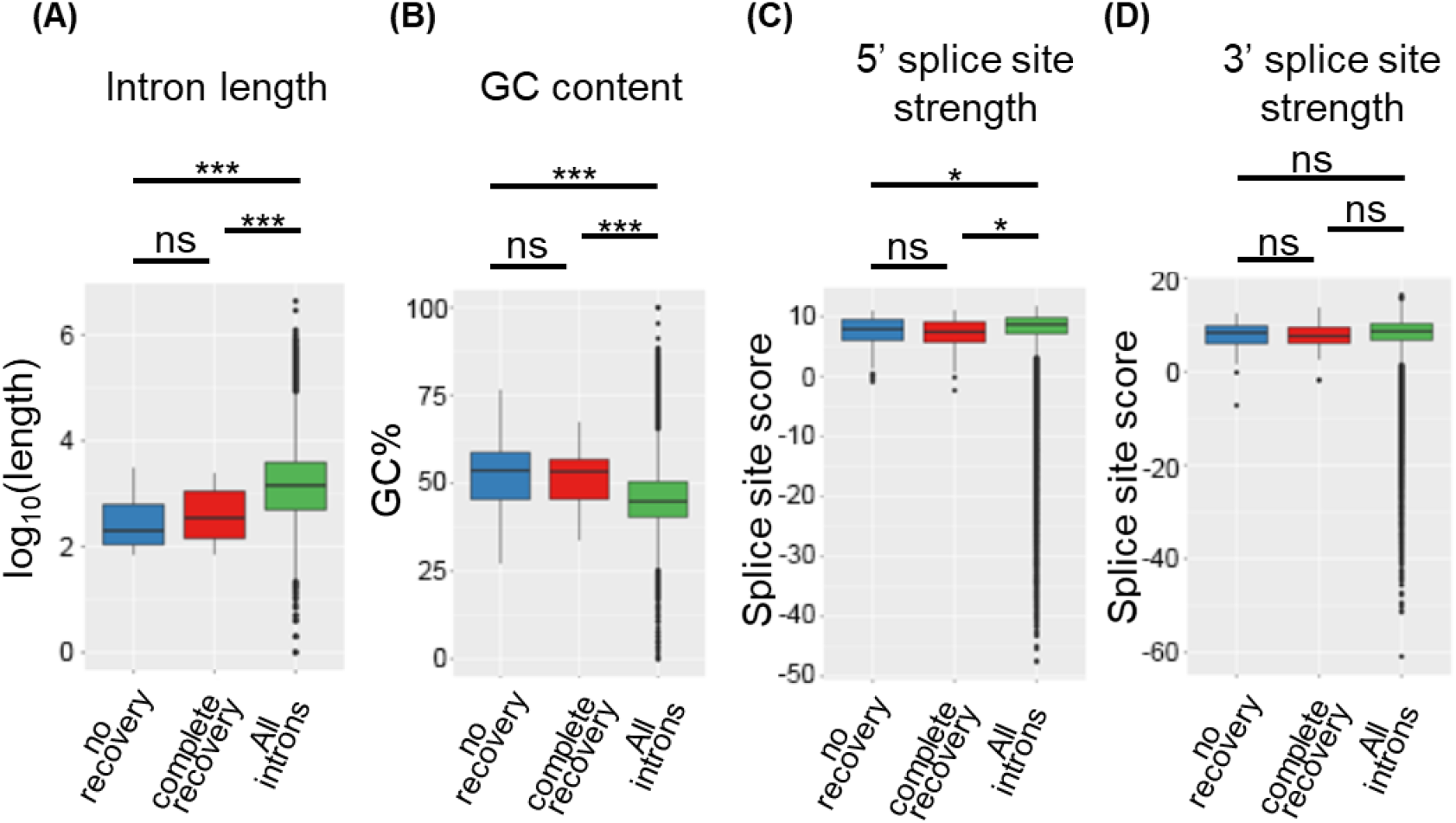
Loci with retained introns have distinguishing characteristics. (A–D) Boxplots showing (A) intron lengths, (B) the GC percentage in intron sequences, and the strength score of the (C) 5’ and (D) 3’ splice sites compared among three groups of introns, namely “no recovery”, “complete recovery”, and all introns from liver-expressed genes (254,005 loci). \**P* < 0.05, \*\*\**P* < 0.001, unpaired Student’s Ltest; ns, not significant.

### 3.8. Validation of IR-recovered genes in the liver by using RT-PCR

Eight genes were chosen for IGV mapping and RT-PCR, which confirmed the recovery of IR upon JTT administration (Figure 5). These genes are involved in RNA splicing (*Ddx5*) and typical liver functions such as glucose and fatty acid metabolic pathways (*Sirt7, Acadm, Decr2*), lipid metabolic processes (*Acadm*), heme biosynthesis and bile acid production (*Cyp27a1, Hsd3b7*), and oxidationreduction processes (*Acadm, Cyp27a1, Hsd3b7*).

**Figure 5.**
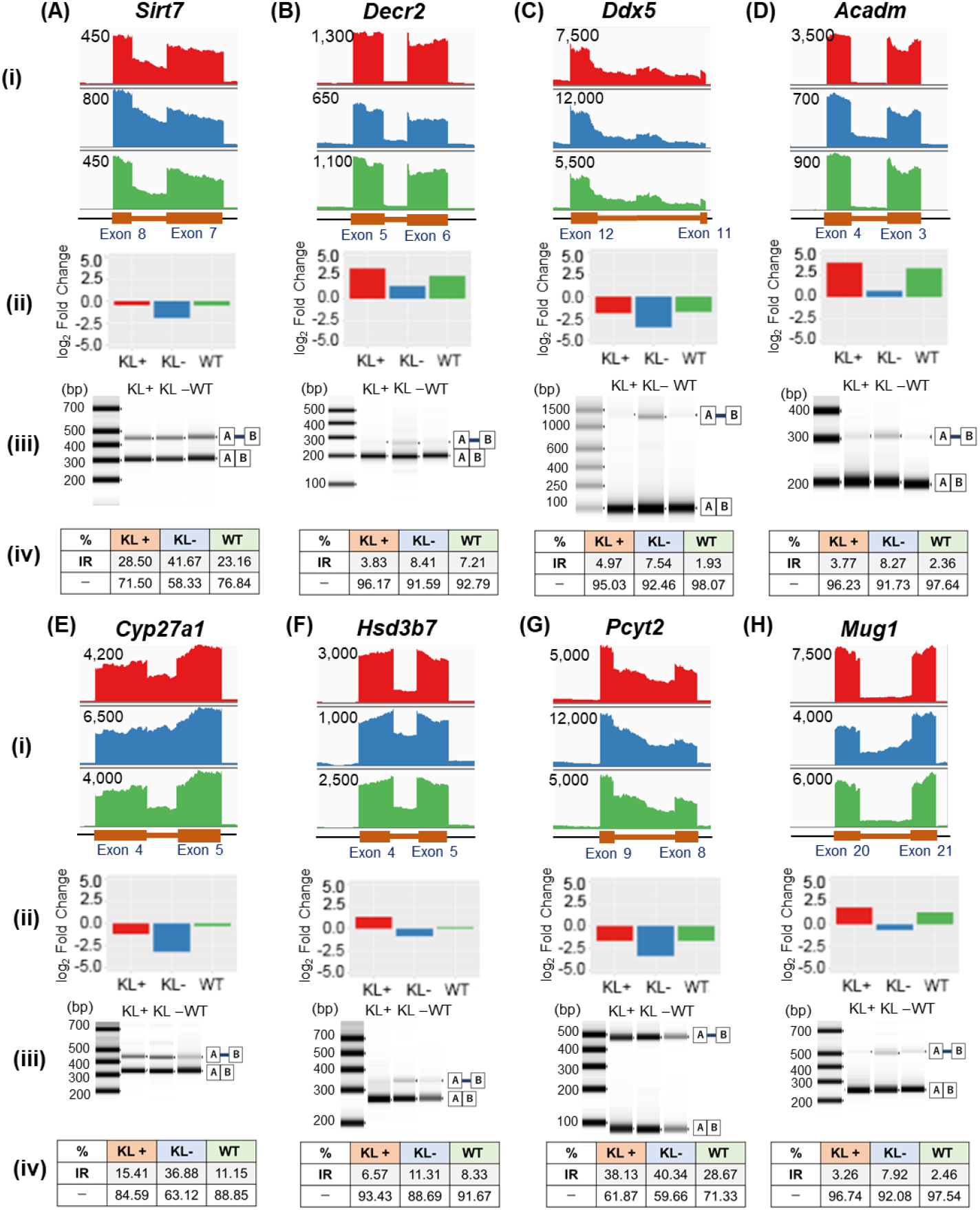
RT-PCR validation of retained introns and their recovery after administration of JTT. (A–H) Eight genes, (A) *Sirt7*, (B) *Decr2*, (C) *Ddx5*, (D) *Acadm*, (E) *Cyp27a1*, (F) *Hsd3b7*, (G) *Pcyt2*, and (H) *Mugl*, were subjected to the following analyses. (i) IGV mapping of reads from KL+, KL−, and WT. The numbers shown to the left of each map represent read counts. (ii) Bar graphs showing fold changes in SJCs relative to IJCs for KL+, KL−, and WT. (iii) RT-PCR validation of RNA expression from KL+, KL−, and WT. A and B indicate exons, with the intervening intron indicated for the higher-molecular-weight product. (iv) The ratio of each transcript as determined using TapeStation. Data from KL+, KL−, and WT are shown in red, blue, and green, respectively.

### 3.9. Biological function of IR-recovered genes in the liver

Possible effects of JTT on the selection of liver retained-intron loci for recovery during premature aging were analyzed (Figure 6). The results obtained from the GO analysis using Biological Process terms in Figure 6A & 6B suggest two conclusions. First, in the comparison between DecIR type (132 loci, 122 genes) of KL+ vs. KL– and IncIR type (190 loci, 179 genes) of KL– vs. WT, genes involved in RNA splicing and mRNA processing are susceptible to IR during the aging process, and these retained introns tend to be recovered by JTT (i.e., the red and blue bars are comparable; Figure 6A). The tendency of genes involved in these processes to be susceptible to IR during aging is consistent with a previous study (Zane et al., 2014). These splicing-related genes can control the expression of downstream genes through the regulation of their alternative splicing. We defined 250 genes as potential splicing-related genes to highlight their importance in the regulation of alternative splicing (see Methods for the definition of splicing-related genes). They included almost all the spliceosomal proteins and spliceosome regulatory proteins. Among 368 gene loci that exhibited a difference in alternative splicing for the comparison of KL+ and KL− in the liver (Table 1-2), 19 genes (data not shown) were detected among the list of 250 potential splicing-related genes. Among these 19 genes, 9 potential splicing-related genes in the liver were subjected to IR, all of which exhibited the recovery pattern in terms of the comparison of the FC among KL+, KL−, and WT (Table 2). Accordingly, JTT appears to work in the direction of recovery, especially for potential splicing-related genes.

**Figure 6.**
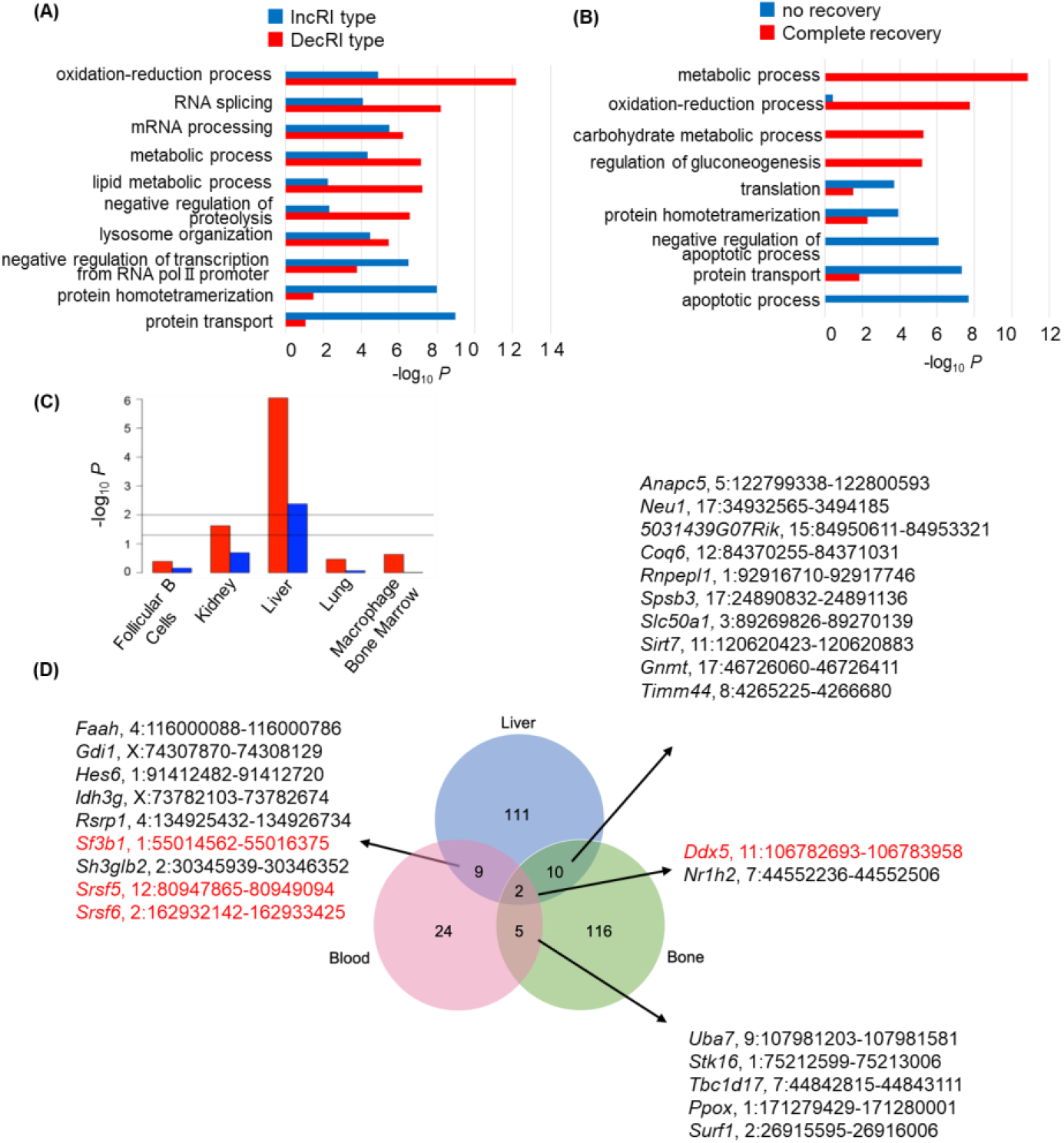
Characterization of genes with recovered retained introns. (A) Bar graphs showing – log_10_ *P* of GO terms enriched for genes whose IR events were significantly changed in the liver. Red bars show enrichment of GO terms in genes with the DecIR type in KL+ in the KL+ and KL– comparison, and blue bars show enrichment of GO terms in genes with the IncIR type in KL– in the KL– and WT comparison. (B) Bar graph showing –log_10_ *P* of GO terms enriched for genes defined as “complete recovery” (red bars) and “no recovery” (blue bars) loci. (C) Cten analysis for genes expressed in the liver with the DecIR type in red and those with the IncIR type in blue. The vertical axis shows the logarithm of the *P*-values of enrichment calculated by Cten. Horizontal lines indicate *P* = 0.05 (lower line) and 0.01 (upper line). (D) A Venn diagram of the DecIR genes showed that *Ddx5* and *Nr1h2* were shared in common among liver, blood, and bone in the recovery process of IR after administration of JTT. Eleven genes were shared between liver and blood, 12 genes were shared between liver and bone, and 7 genes were shared between blood and bone. The overlapping genes between two or three organs, among which splicing-related genes are shown in red, are indicated.

**Table 2.**
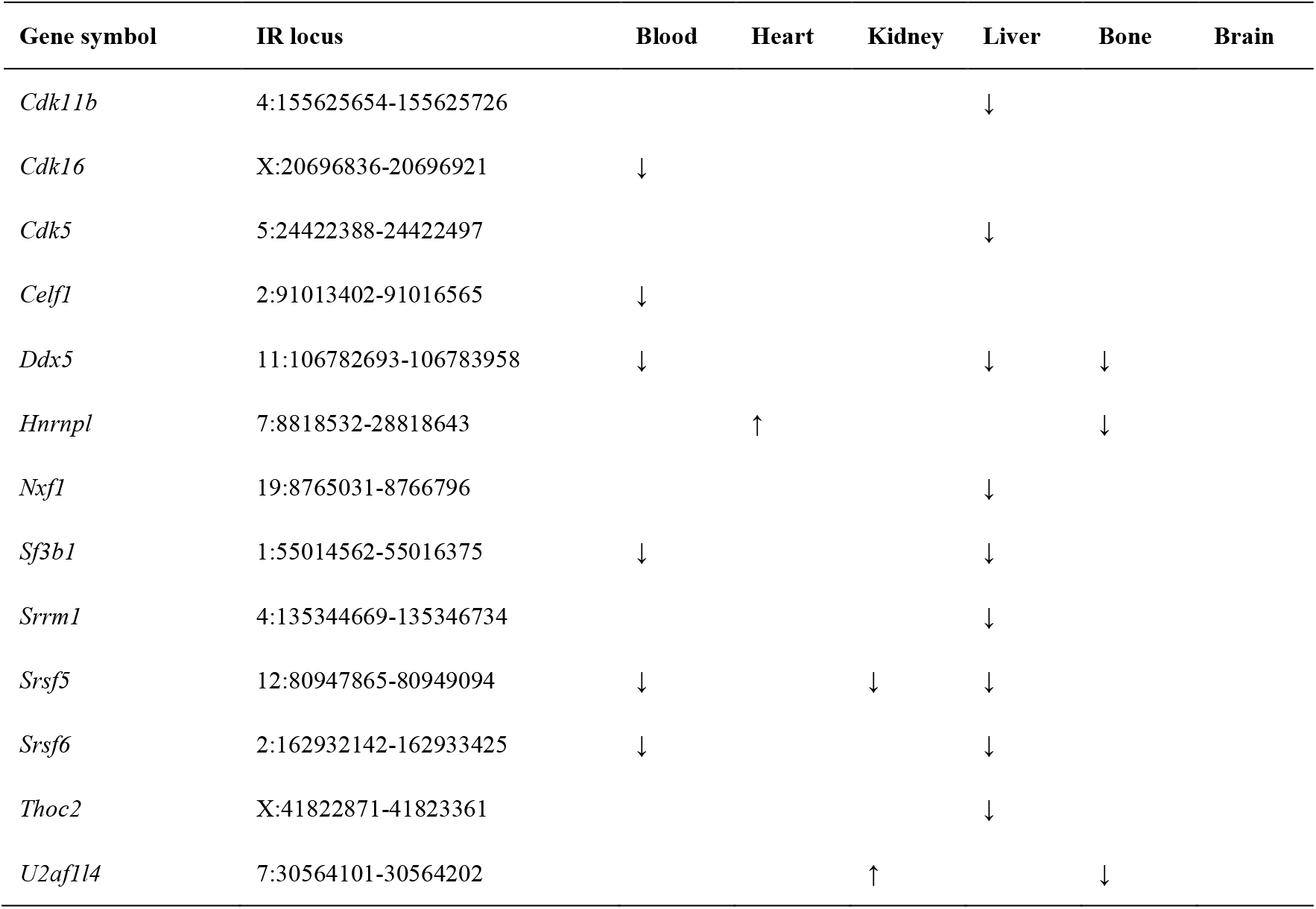
Increase or decrease in the IR ratio of splicing-related genes in blood, heart, kidney, liver, bone, and brain after administration of JTT.

Second, in addition to the recovery of IR for genes related to RNA splicing and RNA processing, IR recovery was not evenly distributed among genes that accumulated retained introns during aging but instead occurred selectively among liver-specific genes. This conclusion was more obvious when loci of “complete recovery” (70 loci) and “no recovery” (120 loci) were compared (Figure 6B). The GO terms such as metabolic process, oxidation-reduction process, carbohydrate metabolic process, and regulation of gluconeogenesis, which represent major functions of the liver, were enriched among genes that underwent “complete recovery” (i.e., red bars), whereas genes in the “no recovery” group are associated with more general functions such as translation, protein transport, and apoptosis (i.e., blue bars). These data suggest that, among genes susceptible to IR during aging, the liver-specific genes selectively underwent a shift in retained introns from the senescent state to the normal state after administration of JTT.

Cten analysis showed that genes with “complete recovery” were significantly enriched among liver-specific genes (Figure 6C; −log_10_ *P* = 6.040), confirming that the IR recovery from premature aging specifically occurred in liver-specific genes.

### 3.10. Some liver metabolic functions were improved to the normal state by administration of JTT

Before discussing a model concerning the mechanism by which only certain genes undergo IR recovery in response to JTT, we highlight the results of 213 genes from the DEG analysis, the expression levels of which were down-regulated in KL− and were recovered to the level of WT in KL+ (Figure 2G). GO analysis of these genes showed that metabolic processes and glucuronidation in the liver likely were up-regulated in KL+ and had recovered after administration of JTT from the down-regulated state in KL– (Figure 2I).

There are many clinical and basic scientific reports on the improvement of liver functions by JTT, typically exemplified by reports that JTT can protect the liver from injury by chemicals administered to patients with cancer or other diseases. A few examples are listed below. JTT protects against isoniazid/rifampicin-induced hepatic injury by modulating oxidative stress and inflammatory responses (Dvinge and Bradley, 2015). JTT exerts protective effects against alcohol-induced liver disease by modulating oxidative stress and the inflammatory response (Monteuuis et al., 2019). JTT has the potential to protect against bromobenzene-induced hepatotoxicity and to modulate oxidative stress (Han et al., 2016). The presence of such a variety of reports convinced us that an improvement in liver function, especially metabolic activities, is one of the major effects of JTT. These published data are consistent with our DEG data described above (Figure 2I).

### 3.11. A model to explain why IR was recovered in a specific subset of genes involving liver metabolism

There were three main differences in the pre-symptomatic state in klotho mice in response to JTT as deduced from the comparison between KL+ and KL–, namely metabolite changes (Figure 1F & S3), changes in the transcriptome as detected by DEGs (Figure 2A-B), and changes in IR (Figure 3). Based on these three fundamental changes, we have proposed a new model to explain why IR was recovered in response to changes in metabolites and the transcriptome. Our model is analogous to the heat shock response reported by Shalgi et al. (2014) As was mentioned in the Introduction, IR is an important mechanism for controlling the rate of protein production based on the retention of intron-containing mRNAs in the nucleus. During the heat shock response in mice, there is widespread retention of introns in transcripts from 1,700 genes, which are enriched for tRNA synthetase, nuclear pore, and spliceosome functions. In contrast, transcripts from 500 genes involving protein-folding functions and oxidation reduction are normally spliced and translated. The transcripts with retained introns are, for the most part, nuclear and untranslated and are mostly spliced post-transcriptionally, in contrast to other normal splicing that occurs co-transcriptionally (Shalgi et al., 2014). These nuclear retained transcripts are presumed to await a signal for splicing that is cued by the release from the stress (Jacob and Smith, 2017). The presence of other similar examples (Boutz et al., 2015; Ninomiya et al., 2011) of stress-induced regulation of IR suggests that this regulatory mechanism is evolutionarily conserved and contributes to the expression of a wide range of genes.

As aging is also a kind of stress, we considered IR during premature aging as analogous to that which occurs during the heat shock response. Figure 7A shows a model for the case of KL−, in which transcripts with retained introns are retained in the nucleus upon the stress signal of aging such that global metabolic activity in liver cells is repressed to save energy. After administration of JTT, as liver-specific metabolic functions are recovered to some extent as described above (Figure 2), transcription-based improvement results in a signal to the liver cells to recover these retained introns by sending a cue for post-transcriptional splicing (Figure 7A(ii)). If this model is correct, we can determine which pathways in the cells are improved by knowing which functions the IR-recovered genes are involved in. In the present case, the recovery of retained introns of genes involved in lipid/glucose metabolic pathways suggests that the liver-specific metabolic functions were recovered. Genes detected by DEG data (Figure 2F & 2G), the Cten analysis (Figure 2H), and its GO analysis (Figure 2I) confirmed that genes involved in metabolic processes were transcriptionally recovered. Also, the recovery of retained introns of genes involved in splicing pathways (Figure 6A, Table 2) suggests that splicing functions were recovered, at least in part.

**Figure 7.**
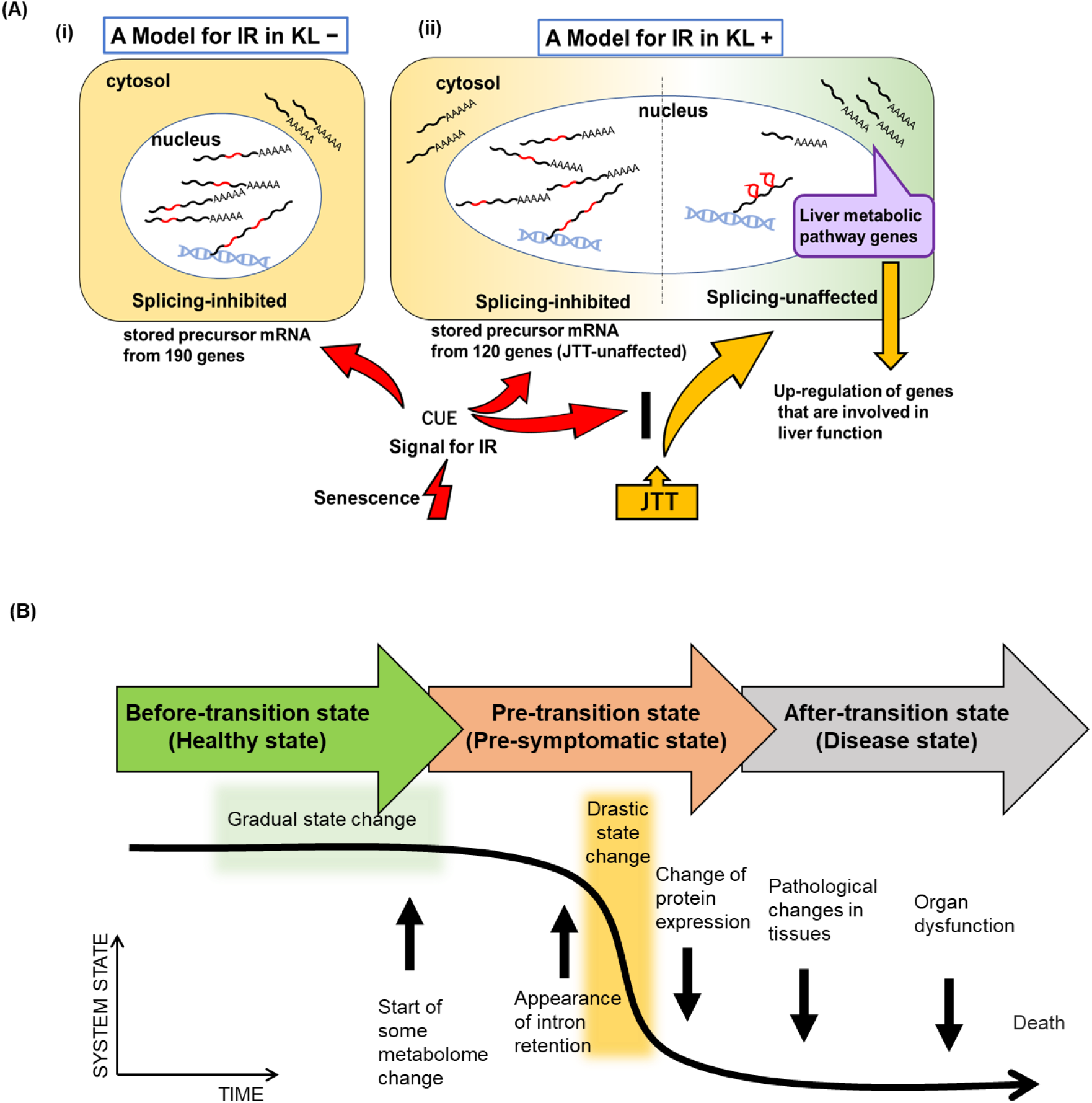
A model to explain why a specific subset of genes with IR was recovered in KL+ by administration of JTT. (A) (i) In the case of KL−, it is presumed that ~200 pre-mRNAs containing retained introns were stored in the nucleus because of the stress signal induced by aging so that the cells can save energy. (ii) In the case of KL+, as liver metabolic stress was recovered after JTT administration, the improved condition of the liver cells led to enhanced post-transcriptional splicing among pre-mRNAs with retained introns involved in metabolism. (B) A schematic representation of the aging process from the healthy state to the disease state.

### 3.12. Overlapping IR-recovered genes among liver, blood, and bone

To examine the characteristics of recovered genes among liver, blood, and bone, DecIR loci from the comparison between KL+ and KL− in liver (132 loci), blood (40 loci), and bone (133 loci) (Figure 3A) were analyzed for overlaps. Figure 6D shows the Venn diagram of this result. It is very interesting to note that the two genes *Ddx5* and *Nr1h2* (LXRβ) were susceptible to undergoing IR recovery in each of these three different organs. These data are indicative of the importance of these two genes in the recovery process of IR by JTT. In particular, IR recovery of *Ddx5* represents the importance of genes involved in RNA splicing in multiple organs during the recovery process by JTT. In the case of 11 genes that underwent IR recovery in both the liver and blood (Figure 6D), four splicing-related genes (*Sf3b1, Ddx5, Srsf5*, and *Srsf6*) were included, again highlighting the importance of these splicing-related genes (Table 2).

IR recovery of *Nr1h2* (LXRβ) probably represents the importance of the metabolic functions of this gene, which apparently functions in the liver, blood, and bone during the recovery process associated with JTT. *Nr1h2* (LXRβ) is a key regulator gene in macrophage function (Rui, 2014). Currently, it is known that LXRs have three major functions. First, LXRs contribute to maintaining cellular cholesterol homeostasis by controlling transcriptional programs involved in lipid homeostasis (Zelcer and Tontonoz, 2006). Macrophages take up oxidized-LDL, and its increased cellular concentration activates LXRs, which induce the efflux of intracellular cholesterols from macrophages to serum apolipoproteins in the form of HDL, which then returns to the liver in a process termed reverse cholesterol transport. This process prevents the contribution of foam-transformed macrophages loaded with cholesterol to atherosclerosis at the inner wall of blood vessels (Zelcer and Tontonoz, 2006). Second, LXRs exhibit anti-inflammatory activity by inhibition of the NFκB– dependent induction of inflammatory gene expression through immune-related stimulation such as by TNF-α or IL-1β (Zelcer and Tontonoz, 2006). Third, LXR plays an important role in the activation of phagocytosis, which is a major function of macrophages. LXR transactivates Mer receptor tyrosine kinase (MERTK), which is a phagocytic receptor, thereby increasing the efficiency of phagocytosis (Kidani and Bensinger, 2012).

Oral administration of JTT can elevate the phagocytic activity of macrophage-like cells in various tissues (Saiki, 2000; Takeno et al., 2015). Similarly, the phytoestrogen flavonoids found in Glycyrrhizae radix, one of the components of JTT, can enhance phagocytosis in a murine macrophage cell line as we described previously (Kaneko et al., 2017). JTT contains various types of flavonoids, some of which have been identified as selective agonists of LXR, according to molecular-docking analysis *in silico* (Fouache et al., 2019). Moreover, a representative phytoestrogen has been reported to alter the LXR signaling system *in vivo* and *in vitro* (Luo et al., 2018). Because JTT reduced the age-related accumulation of IR in *Nr1h2* in klotho mice, it could help to maintain a healthy immune system in elderly individuals by influencing macrophage functions.

Among the 11 IR-recovered loci that overlapped between liver and blood (Figure 6D), *Faah* is connected with macrophage function through lipid metabolism (Yoshioka et al., 2019), and *Gdi1* is connected with interferon-γ activity through innate immunity (Fukaya et al., 2018). Among 12 IR-recovered loci that overlapped between liver and bone (Figure 6D), *Sirt1* is positively involved in macrophage self-renewal activity, during which it functions to increase their proliferative capacity during bone marrow-derived macrophage differentiation (Imperatore et al., 2017). The overlap of such IR-recovered genes among these three organs might indicate a contribution of the systemic efficacy of JTT through IR recovery of these genes in connection with that of *Nr1h2*.

### 3.13. The mechanism for aging-related IR in transcripts and possible gene specificity of JTT-related recovery

Regarding the mechanisms through which transcripts retained their introns and by which a specific group of transcripts was recovered, we suggest that IR-susceptible genes during aging and IR-recovered genes after JTT administration are transcriptionally controlled by a very few TFs that are specific to the liver, bone, and blood (Figure 8 & S6). We first examined enrichment for the binding sites of TFs in the IR-recovered genes in the liver (122 genes, 132 loci), in the blood (38 genes, 40 loci), and in the bone (119 genes, 133 loci) that were categorized as DecIR (KL+ vs KL−) (Figure 8A). These data clearly showed that different sets of TFs were used for transcription in IR-recovered genes in these organs. This observation prompted us to examine how many IR-increased genes or IR-recovered genes in the respective organs have a common TF-binding site in the promoter region spanning 500 bases upstream from the transcription start site. In the case of a decrease in retained introns in response to administration of JTT, 121 of 122 genes (132 loci) in the liver had binding sites for at least one of four TFs, namely AR-halfsite, Tgif2, KLF14, and Hoxa11, as shown in Figure 8B. Similarly, in the blood, 38 of 38 genes (40 loci) had binding sites for at least one of four TFs, namely PU.1:IRF8, EIK1, Hoxa13, and Sp2. In the bone, 114 of 119 genes (133 loci) had binding sites for at least one of four TFs, namely Sp2, SCL, GABPA, and Nanog. These data clearly show that a distinct set of genes sharing the same TF-binding sites was affected during the IR-increasing process of aging and during the IR-decreasing process after administration of JTT and were networked.

**Figure 8.**
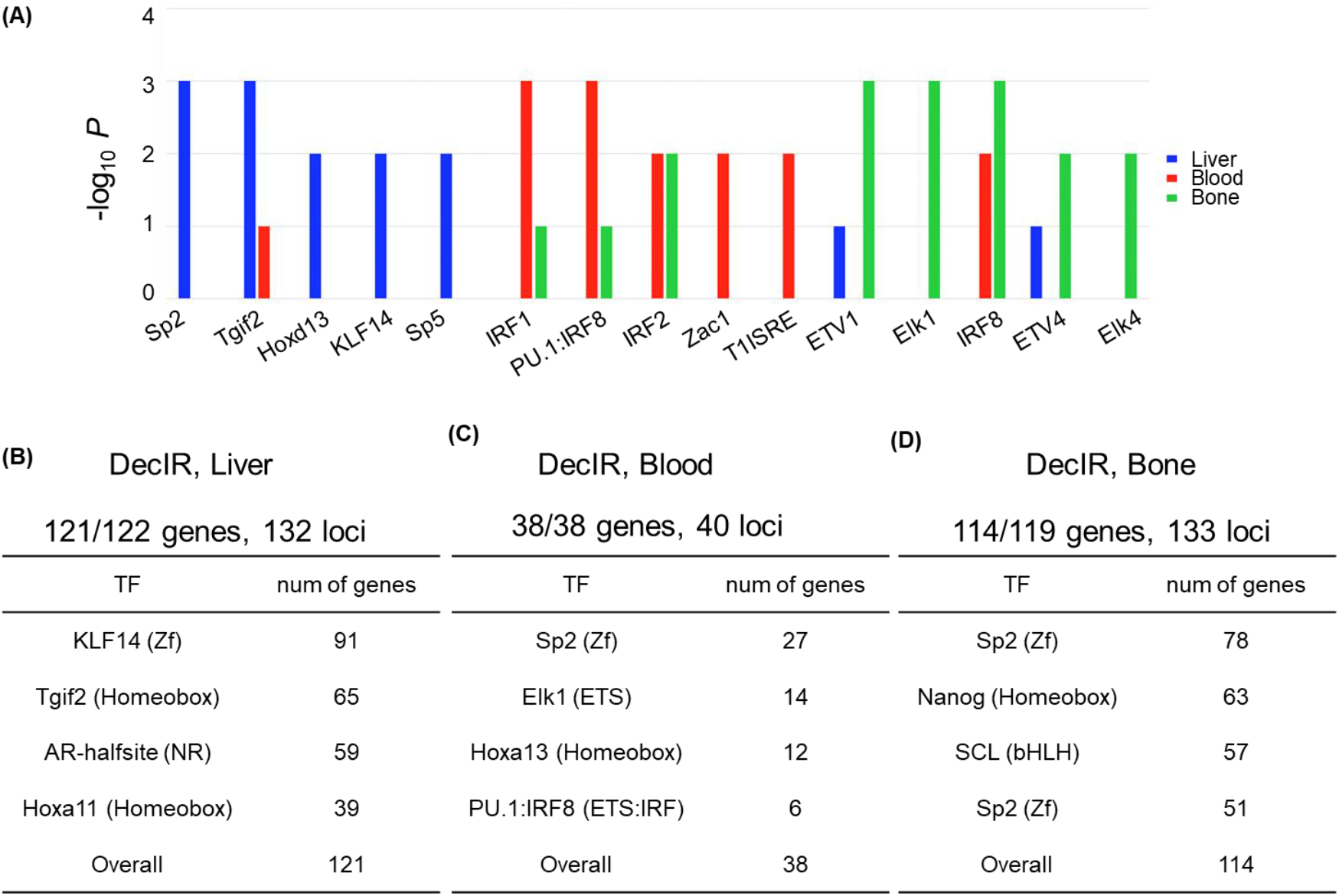
A very few TFs may control genes with DecIR. (A) Bar graph showing TF binding motifs that are significantly enriched in genes with DecIR from blood, liver, and bone. (The vertical axis shows the negative log-transformed *P*-value for enrichment.) (B-D) Those TF-binding motifs that are more prevalent among affected genes in the liver (B), blood (C), and bone (D) are shown.

Similar analyses were performed on the IR-increased genes in the liver (179 genes, 190 loci), in the blood (14 genes, 15 loci), and in the bone (77 genes, 82 loci) that were categorized as IncIR (KL− vs. WT−) (Figure S7A–B). This type of analysis suggests which TFs may have been involved in the generation of retained introns that increased during progression of the pre-disease process. Most TFs were organ specific, as only a few TFs were shared among liver, blood, and bone as shown in Figure S6. All of the TFs listed in Figure S7, which are listed in Figure S7, might play a more general role in aging.

There is emerging evidence that mRNA splicing is controlled by multi-layered mechanisms that involve transcription and epigenetics (Braunschweig et al., 2014; Lev Maor et al., 2015; Luco et al., 2011). Therefore, it is interesting to consider the possibility that TFs, chromatin constituents, and epigenetic factors such as those related to histone modification and DNA methylation not only control transcription but also regulate splicing (Lev Maor et al., 2015; Luco et al., 2011). These factors can affect the rate of RNA polymerase II elongation, which in turn affects the pattern of alternative splicing. Accordingly, it is possible that accumulation of retained introns during aging in klotho mice was caused by a change in the elongation rate of transcription. As shown in Figure S7, there are several TF-binding sites that are enriched in the genes that were susceptible to IR during aging in liver (190 loci, 179 genes), in blood (15 loci, 14 genes), and in bone (82 loci, 77 genes). Accordingly, an aging signal such as fasting may lead specific TFs to retard the transcription rate, resulting in IR. Also, it is possible that some active ingredients of JTT in cooperation with these TFs could contribute to the recovery of IR in liver (132 loci, 122 genes), in blood (40 loci, 38 genes), and in bone (133 loci, 119 genes).

Also, it is interesting to note that the concentration of certain nucleotide diphosphates (UDP and GDP) was decreased in KL– and recovered to WT levels in KL+ (Figure S3C). If a low concentration of nucleotides triggers a change in the elongation rate of transcription (Kwak and Lis, 2013), it is also possible that this may have induced the accumulation of retained introns. The recovery of such concentrations to the normal level in KL+ is consistent with the recovery of retained introns in KL+.

Although there are instances in which chemical compounds influence alternative splicing by directly interacting with splicing factors (Kataoka, 2017), there has been no report of changes in alternative splicing patterns that result from an interaction with a TF. It is of interest to examine whether the active ingredients of a Kampo medicine can interact with TFs to influence alternative splicing patterns co-transcriptionally or post-transcriptionally. The current results regarding the reduction of retained introns by JTT will provide a rich resource for the subsequent studies of the regulation of alternative splicing.

### 3.14. Determining whether IR that occurs during klotho mouse aging is observable in other instances such as the aging of normal mice

Based on the data described above about klotho mice, we next asked whether IRs can be observed in the liver of WT mice during the aging process. Figure 9A shows the analysis flow chart for this study (see also Materials and Method section 2.7), in which we first chose 856 loci as possible candidate loci for IR after quality control (i.e., removal of loci with exons with coverage of <5 or intron lengths of <50 bases) among all possible mouse IR loci. We then examined the extent of IR in these genes in WT mice from 2 months to 30 months. Figure 9B(i) and (iii) show the distribution of the number of loci with PSI > 0.1 and the relative abundance of PSI levels for these genes at specific ages, respectively, both of which indicated that there was no tendency toward increasing PSI among the 856 loci in the liver as a whole. As described above in the case of klotho mice, 190 loci increased their introns in the liver during aging and were characterized as IncIR based on the comparison between KL– and WT (Figure 3C). Accordingly, with the possibility that these 190 loci characterized in klotho mice might have also tended to accumulate introns in the WT mice during aging, we checked the distribution of the number of loci with PSI > 0.1 among these 190 loci and their relative abundance by PSI level at the specific ages as shown in Figure 9B(ii) and (iv). Interestingly, based on this selection, we clearly demonstrated that introns accumulated during aging even in the WT mice. In particular, it should be noted that accumulation of introns in these 190 loci was first noted at the time of 15 months (Figure 9B(iv)), less than half of the life-span of WT mice (i.e., about 2.5 years), which corresponds to ~7 weeks of age in klotho mice, which ultimately resulted in a significant difference (*P* = 0.0299) in accumulated introns for the comparison between 2 months and 30 months. These data suggest that genes susceptible to IR during aging are shared to some extent between WT mice and klotho mice and could be evolutionarily conserved.

**Figure 9.**
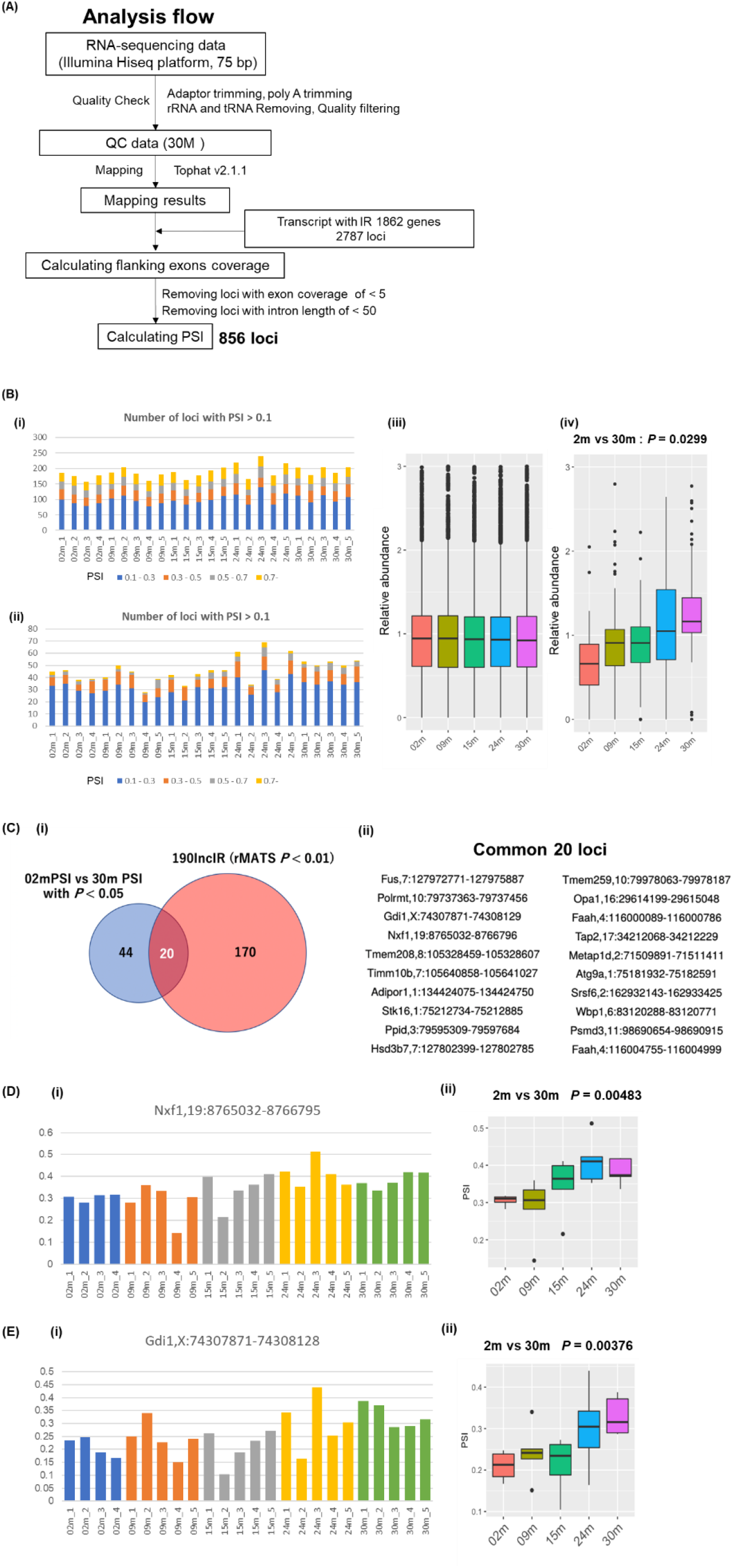
Retained introns were observed at the pre-symptomatic state of aging of WT mice. (A) Flow of analysis of IR among liver mRNAs in WT mice using the PSI calculation method. (This analysis used the GSE75192 dataset downloaded from the NCBI GEO database.) (B) Analyses of the number of loci with PSI > 0.1 during the aging process of WT mice at five different timepoints (2 months, 9 months, 15 months, 24 months, and 30 months). (i) Number of intronic loci with PSI values > 0.1 at 856 loci. (ii) Number of intronic loci with PSI values > 0.1 at 190 loci that were significantly different between KL– and WT. (iii) Boxplot of relative abundance calculated from the average PSI of all individuals per intron locus at 856 loci. (iv) Boxplot of relative abundance calculated from the average PSI of all individuals per intron locus at 190 loci that were significantly different between KL– and WT. (C)(i) Venn diagram comparing 64 loci with increased IR with a significant difference of *P* < 0.05 between 2 months and 30 months of GSE75192 and 190 loci with increased IR with a significant difference of *P* < 0.01 between KL– and WT. (ii) Twenty common loci with increased IR in C(i). (D)(i) Bar graph of PSI values of the intron locus of *Nxf1* with significant differences between 2 months and 30 months. (ii) Boxplot of PSI values at the intron locus of *Nxf1*. (iii) Bar graph of PSI value of the intron locus of *Gdi1* with significant differences between 2 months and 30 months. (ii) Boxplot of PSI value at the intron locus of *Gdi1*.

The following analysis confirmed that WT and klotho mice do have some IR susceptible genes in common. From the analysis of PSI of the WT mice between 2 months and 30 months, we could list 64 loci that showed a significant difference with *P* < 0.05. Remarkably, we demonstrated that 20 loci among these 64 loci were also found among the 190 loci with IncIR based on the comparison between KL− and WT (Figure 9C), confirming the above notion. The aging process differs between WT mice and klotho mice, as the aging of WT mice is due to the gradual deterioration of the molecular components of the cell whereas that of klotho mice is due to the high concentration of phosphorous in the blood and subsequent perturbation of calcium-related metabolism (see Introduction). These 20 overlapping loci should represent common aspects of the aging process in WT mice and klotho mice. Figure 9D-E shows the process of intron accumulation in two particular genes during aging, *Nxf1* and *Gdi1*, each of which was listed as one of the completely recovered loci in klotho mice (70 loci; Figure 3C) although each began to accumulate introns at different times, namely, the former at 15 months (Figure 9D(ii)) and the latter at 24 months (Figure 9E(ii)) in WT mice.

### 3.15. Future Perspectives

The present study provides evidence that IR may be an evolutionarily conserved mechanism and can be observed at the pre-symptomatic state during aging. Accordingly, it represents a new marker for this state. This study is also the first to shed light on the comprehensive mechanism of a multi-herbal medicine from the viewpoint of systems biology. It is interesting to note that the ancient Chinese medical textbook *Inner Canon of the Yellow Emperor* stated that the pre-disease state should be treated early with Kampo medicine (UNESCO, 2011). The present data provide a molecular basis for this historical statement. Kampo medicine is a precious heritage in Japan, and there are similar medicines used around the world such as traditional Chinese medicines (Patwardhan et al., 2005), traditional medicines in India (Ayurveda) (Patwardhan et al., 2005), and ancient Greek medicines (Unani medicine). To elucidate the mechanisms of these medicines, which are the result of human wisdom that has accumulated over several thousand years, and to pass on this knowledge to subsequent generations are the main goals of our research.

## Availability

The original RNA-seq datasets have been deposited in the DDBJ Sequence Read Archive under accession numbers DRR167982–DRR167990, which are linked to the BioProject accession number PRJDB7898.

## Acknowledgements

The present work was financially supported by the Foundation for Advancement of International Science and Zenick Co. represented by Mr. Y. Otake and Mr. T. Sugino, respectively. We thank Mr. K. Onodera for management and discussions and Ms. A. Koizumi for making the illustrations.

## References

Adusumalli, S., Ngian, Z.K., Lin, W.Q., Benoukraf, T., and Ong, C.T. (2019). Increased intron retention is a post-transcriptional signature associated with progressive aging and Alzheimer’s disease. Aging Cell 18(3), e12928. DOI:10.1111/acel.12928.

Aramillo Irizar, P., Schäuble, S., Esser, D., Groth, M., Frahm, C., Priebe, S., Baumgart, M., Hartmann, N., Marthandan, S., Menzel, U., et al. (2018). Transcriptomic alterations during ageing reflect the shift from cancer to degenerative diseases in the elderly. Nature Communications 9(1), 327. DOI:10.1038/s41467-017-02395-2.

Bolger, A.M., Lohse, M., and Usadel, B. (2014). Trimmomatic: a flexible trimmer for Illumina sequence data. Bioinformatics 30(15), 2114–2120. DOI:10.1093/bioinformatics/btu170.

Boutz, P.L., Bhutkar, A., and Sharp, P.A. (2015). Detained introns are a novel, widespread class of post-transcriptionally spliced introns. Genes Dev 29(1), 63–80. DOI:10.1101/gad.247361.114.

Braunschweig, U., Barbosa-Morais, N.L., Pan, Q., Nachman, E.N., Alipanahi, B., Gonatopoulos-Pournatzis, T., Frey, B., Irimia, M., and Blencowe, B.J. (2014). Widespread intron retention in mammals functionally tunes transcriptomes. Genome Res 24(11), 1774–1786. DOI:10.1101/gr.177790.114.

Deschenes, M., and Chabot, B. (2017). The emerging role of alternative splicing in senescence and aging. Aging Cell 16(5), 918–933. DOI:10.1111/acel.12646.

Desvergne, B., Michalik, L., and Wahli, W. (2006). Transcriptional regulation of metabolism. Physiol Rev 86(2), 465–514. DOI:10.1152/physrev.00025.2005.

Dvinge, H., and Bradley, R.K. (2015). Widespread intron retention diversifies most cancer transcriptomes. Genome Med 7(1), 45. DOI:10.1186/s13073-015-0168-9.

Fouache, A., Zabaiou, N., De Joussineau, C., Morel, L., Silvente-Poirot, S., Namsi, A., Lizard, G., Poirot, M., Makishima, M., Baron, S., et al. (2019). Flavonoids differentially modulate liver X receptors activity-Structure-function relationship analysis. J Steroid Biochem Mol Biol 190, 173–182. DOI:10.1016/j.jsbmb.2019.03.028.

Fukao, T., Mitchell, G., Sass, J.O., Hori, T., Orii, K., and Aoyama, Y. (2014). Ketone body metabolism and its defects. J Inherit Metab Dis 37(4), 541–551. DOI:10.1007/s10545-014-9704-9.

Fukaya, S., Nagatsu, A., and Yoshioka, H. (2018). The Kampo formula “Juzen-taiho-to” exerts protective effects on ethanol-induced liver injury mice. Fundamental Toxicological Sciences 5(3), 105–112. DOI:DOI:10.2131/fts.5.105

Giudice, G., Sanchez-Cabo, F., Torroja, C., and Lara-Pezzi, E. (2016). ATtRACT-a database of RNA-binding proteins and associated motifs. Database (Oxford) 2016. DOI:10.1093/database/baw035.

Goswami, C., Dezaki, K., Wang, L., Inui, A., Seino, Y., and Yada, T. (2019). Ninjin-yoeito activates ghrelin-responsive and unresponsive NPY neurons in the arcuate nucleus and counteracts cisplatin-induced anorexia. Neuropeptides 75, 58–64. DOI:10.1016/j.npep.2019.03.001.

Gunewardena, S.S., Yoo, B., Peng, L., Lu, H., Zhong, X., Klaassen, C.D., and Cui, J.Y. (2015). Deciphering the Developmental Dynamics of the Mouse Liver Transcriptome. PLoS One 10(10), e0141220. DOI:10.1371/journal.pone.0141220.

Han, H., Braunschweig, U., Gonatopoulos-Pournatzis, T., Weatheritt, R.J., Hirsch, C.L., Ha, K.C.H., Radovani, E., Nabeel-Shah, S., Sterne-Weiler, T., Wang, J., et al. (2017). Multilayered Control of Alternative Splicing Regulatory Networks by Transcription Factors. Mol Cell 65(3), 539–553 e537. DOI:10.1016/j.molcel.2017.01.011.

Han, H.S., Kang, G., Kim, J.S., Choi, B.H., and Koo, S.H. (2016). Regulation of glucose metabolism from a liver-centric perspective. Exp Mol Med 48, e218. DOI:10.1038/emm.2015.122.

Heinz, S., Benner, C., Spann, N., Bertolino, E., Lin, Y.C., Laslo, P., Cheng, J.X., Murre, C., Singh, H., and Glass, C.K. (2010). Simple combinations of lineage-determining transcription factors prime cis-regulatory elements required for macrophage and B cell identities. Mol Cell 38(4), 576–589. DOI:10.1016/j.molcel.2010.05.004.

Huang da, W., Sherman, B.T., and Lempicki, R.A. (2009). Systematic and integrative analysis of large gene lists using DAVID bioinformatics resources. Nat Protoc 4(1), 44–57. DOI:10.1038/nprot.2008.211.

Imperatore, F., Maurizio, J., Vargas Aguilar, S., Busch, C.J., Favret, J., Kowenz-Leutz, E., Cathou, W., Gentek, R., Perrin, P., Leutz, A., et al. (2017). SIRT1 regulates macrophage self-renewal. Embo j 36(16), 2353–2372. DOI:10.15252/embj.201695737.

Jacob, A.G., and Smith, C.W.J. (2017). Intron retention as a component of regulated gene expression programs. Hum Genet 136(9), 1043–1057. DOI:10.1007/s00439-017-1791-x.

Kadota, Y., Jam, F.A., Yukiue, H., Terakado, I., Morimune, T., Tano, A., Tanaka, Y., Akahane, S., Fukumura, M., Tooyama, I., et al. (2020). Srsf7 Establishes the Juvenile Transcriptome through Age-Dependent Alternative Splicing in Mice. iScience 23(3), 100929. DOI:10.1016/j.isci.2020.100929.

Kaneko, A., Matsumoto, T., Matsubara, Y., Sekiguchi, K., Koseki, J., Yakabe, R., Aoki, K., Aiba, S., and Yamasaki, K. (2017). Glucuronides of phytoestrogen flavonoid enhance macrophage function via conversion to aglycones by beta-glucuronidase in macrophages. Immun Inflamm Dis 5(3), 265–279. DOI:10.1002/iid3.163.

Kataoka, N. (2017). Modulation of aberrant splicing in human RNA diseases by chemical compounds. Hum Genet 136(9), 1237–1245. DOI:10.1007/s00439-017-1789-4.

Kidani, Y., and Bensinger, S.J. (2012). Liver X receptor and peroxisome proliferator-activated receptor as integrators of lipid homeostasis and immunity. Immunol Rev 249(1), 72–83. DOI:10.1111/j.1600-065X.2012.01153.x.

Kuro-o, M., Matsumura, Y., Aizawa, H., Kawaguchi, H., Suga, T., Utsugi, T., Ohyama, Y., Kurabayashi, M., Kaname, T., Kume, E., et al. (1997). Mutation of the mouse klotho gene leads to a syndrome resembling ageing. Nature 390(6655), 45–51. DOI:10.1038/36285.

Kuro, O.M. (2018). Molecular Mechanisms Underlying Accelerated Aging by Defects in the FGF23-Klotho System. Int J Nephrol 2018, 9679841. DOI:10.1155/2018/9679841.

Kwak, H., and Lis, J.T. (2013). Control of transcriptional elongation. Annu Rev Genet 47, 483–508. DOI:10.1146/annurev-genet-110711-155440.

Langmead, B., and Salzberg, S.L. (2012). Fast gapped-read alignment with Bowtie 2. Nat Methods 9(4), 357–359. DOI:10.1038/nmeth.1923.

Lejeune, F., and Maquat, L.E. (2005). Mechanistic links between nonsense-mediated mRNA decay and pre-mRNA splicing in mammalian cells. Curr Opin Cell Biol 17(3), 309–315. DOI:10.1016/j.ceb.2005.03.002.

Lev Maor, G., Yearim, A., and Ast, G. (2015). The alternative role of DNA methylation in splicing regulation. Trends Genet 31(5), 274–280. DOI:10.1016/j.tig.2015.03.002.

Liu, Y., Gonzalez-Porta, M., Santos, S., Brazma, A., Marioni, J.C., Aebersold, R., Venkitaraman, A.R., and Wickramasinghe, V.O. (2017). Impact of Alternative Splicing on the Human Proteome. Cell Rep 20(5), 1229–1241. DOI:10.1016/j.celrep.2017.07.025.

Lopez-Otin, C., Blasco, M.A., Partridge, L., Serrano, M., and Kroemer, G. (2013). The hallmarks of aging. Cell 153(6), 1194–1217. DOI:10.1016/j.cell.2013.05.039.

Luco, R.F., Allo, M., Schor, I.E., Kornblihtt, A.R., and Misteli, T. (2011). Epigenetics in alternative pre-mRNA splicing. Cell 144(1), 16–26. DOI:10.1016/j.cell.2010.11.056.

Luo, T., Miranda-Garcia, O., Sasaki, G., Wang, J., and Shay, N.F. (2018). Genistein and daidzein decrease food intake and body weight gain in mice, and alter LXR signaling in vivo and in vitro. Food Funct 9(12), 6257–6267. DOI:10.1039/c8fo01718b.

Maruyama, N., Ishigami, A., and Kondo, Y. (2010). Pathophysiological significance of senescence marker protein-30. Geriatr Gerontol Int 10 Suppl 1, S88–98. DOI:10.1111/j.1447-0594.2010.00586.x.

Matsumoto, T., Sakurai, M.H., Kiyohara, H., and Yamada, H. (2000). Orally administered decoction of Kampo (Japanese herbal) medicine, “Juzen-Taiho-To” modulates cytokine secretion and induces NKT cells in mouse liver. Immunopharmacology 46(2), 149–161. DOI:10.1016/S0162-3109(99)00166-6.

Mayeda, A., Munroe, S.H., Caceres, J.F., and Krainer, A.R. (1994). Function of conserved domains of hnRNP A1 and other hnRNP A/B proteins. EMBO J 13(22), 5483–5495. DOI:10.1002/j.1460-2075.1994.tb06883.x

Monteuuis, G., Wong, J.J.L., Bailey, C.G., Schmitz, U., and Rasko, J.E.J. (2019). The changing paradigm of intron retention: regulation, ramifications and recipes. Nucleic Acids Res 47(22), 11497–11513. DOI:10.1093/nar/gkz1068.

Nabeshima, Y. (2002). Klotho: a fundamental regulator of aging. Ageing Res Rev 1(4), 627–638. DOI:10.1016/s1568-1637(02)00027-2.

Naro, C., Jolly, A., Di Persio, S., Bielli, P., Setterblad, N., Alberdi, A.J., Vicini, E., Geremia, R., De la Grange, P., and Sette, C. (2017). An Orchestrated Intron Retention Program in Meiosis Controls Timely Usage of Transcripts during Germ Cell Differentiation. Dev Cell 41(1), 82–93 e84. DOI:10.1016/j.devcel.2017.03.003.

Niemela, E.H., Oghabian, A., Staals, R.H., Greco, D., Pruijn, G.J., and Frilander, M.J. (2014). Global analysis of the nuclear processing of transcripts with unspliced U12-type introns by the exosome. Nucleic Acids Res 42(11), 7358–7369. DOI:10.1093/nar/gku391.

Ninomiya, K., Kataoka, N., and Hagiwara, M. (2011). Stress-responsive maturation of Clk1/4 pre-mRNAs promotes phosphorylation of SR splicing factor. J Cell Biol 195(1), 27–40. DOI:10.1083/jcb.201107093.

Onishi, Y., Yamaura, T., Tauchi, K., Sakamoto, T., Tsukada, K., Nunome, S., Komatsu, Y., and Saiki, I. (1998). Expression of the anti-metastatic effect induced by Juzen-taiho-to is based on the content of Shimotsu-to constituents. Biol Pharm Bull 21(7), 761–765. DOI:10.1248/bpb.21.761.

Padgett, R.A., Grabowski, P.J., Konarska, M.M., Seiler, S., and Sharp, P.A. (1986). Splicing of messenger RNA precursors. Annu Rev Biochem 55, 1119–1150. DOI:10.1146/annurev.bi.55.070186.005351.

Patel, A.A., and Steitz, J.A. (2003). Splicing double: insights from the second spliceosome. Nat Rev Mol Cell Biol 4(12), 960–970. DOI:10.1038/nrm1259.

Patwardhan, B., Warude, D., Pushpangadan, P., and Bhatt, N. (2005). Ayurveda and traditional Chinese medicine: a comparative overview. Evid Based Complement Alternat Med 2(4), 465–473. DOI:10.1093/ecam/neh140.

Pimentel, H., Parra, M., Gee, S.L., Mohandas, N., Pachter, L., and Conboy, J.G. (2016). A dynamic intron retention program enriched in RNA processing genes regulates gene expression during terminal erythropoiesis. Nucleic Acids Res 44(2), 838–851. DOI:10.1093/nar/gkv1168.

Razzaque, M.S. (2009). The FGF23-Klotho axis: endocrine regulation of phosphate homeostasis. Nat Rev Endocrinol 5(11), 611–619. DOI:10.1038/nrendo.2009.196.

Robinson, J.T., Thorvaldsdottir, H., Winckler, W., Guttman, M., Lander, E.S., Getz, G., and Mesirov, J.P. (2011). Integrative genomics viewer. Nat Biotechnol 29(1), 24–26. DOI:10.1038/nbt.1754.

Robinson, M.D., McCarthy, D.J., and Smyth, G.K. (2010). edgeR: a Bioconductor package for differential expression analysis of digital gene expression data. Bioinformatics 26(1), 139–140. DOI:10.1093/bioinformatics/btp616.

Rui, L. (2014). Energy metabolism in the liver. Compr Physiol 4(1), 177–197. DOI:10.1002/cphy.c130024.

Saiki, I. (2000). A Kampo medicine “Juzen-taiho-to”--prevention of malignant progression and metastasis of tumor cells and the mechanism of action. Biol Pharm Bull 23(6), 677–688. DOI:10.1248/bpb.23.677.

Shalgi, R., Hurt, J.A., Lindquist, S., and Burge, C.B. (2014). Widespread inhibition of posttranscriptional splicing shapes the cellular transcriptome following heat shock. Cell Rep 7(5), 1362–1370. DOI:10.1016/j.celrep.2014.04.044.

Shen, S., Park, J.W., Lu, Z.X., Lin, L., Henry, M.D., Wu, Y.N., Zhou, Q., and Xing, Y. (2014). rMATS: robust and flexible detection of differential alternative splicing from replicate RNA-Seq data. Proc Natl Acad Sci U S A 111(51), E5593–5601. DOI:10.1073/pnas.1419161111.

Shimada, T., Kudo, T., Akase, T., and Aburada, M. (2008). Preventive effects of Bofutsushosan on obesity and various metabolic disorders. Biol Pharm Bull 31(7), 1362–1367. DOI:10.1248/bpb.31.1362.

Shoemaker, J.E., Lopes, T.J.S., Ghosh, S., Matsuoka, Y., Kawaoka, Y., and Kitano, H. (2012). CTen: a web-based platform for identifying enriched cell types from heterogeneous microarray data. BMC Genomics 13(1), 460. DOI:10.1186/1471-2164-13-460.

Solana, R., Pawelec, G., and Tarazona, R. (2006). Aging and innate immunity. Immunity 24(5), 491–494. DOI:10.1016/j.immuni.2006.05.003.

Stegeman, R., and Weake, V.M. (2017). Transcriptional Signatures of Aging. J Mol Biol 429(16), 2427–2437. DOI:10.1016/j.jmb.2017.06.019.

Takeno, N., Inujima, A., Shinohara, K., Yamada, M., Shibahara, N., Sakurai, H., Saiki, I., and Koizumi, K. (2015). Immune adjuvant effect of Juzentaihoto, a Japanese traditional herbal medicine, on tumor vaccine therapy in a mouse model. Int J Oncol 47(6), 2115–2122. DOI:10.3892/ijo.2015.3208.

Trapnell, C., Pachter, L., and Salzberg, S.L. (2009). TopHat: discovering splice junctions with RNA-Seq. Bioinformatics 25(9), 1105–1111. DOI:10.1093/bioinformatics/btp120.

Ullrich, S., and Guigo, R. (2019). Dynamic changes in intron retention are tightly associated with regulation of splicing factors and proliferative activity during B-cell development. Nucleic Acids Res. DOI:10.1093/nar/gkz1180.

UNESCO (2011). Huang Di Nei Jing (Yellow Emperor’s Inner Canon) (Beijing: National Library of China).

Vu, T.D., Iwasaki, Y., Shigenobu, S., Maruko, A., Oshima, K., Iioka, E., Huang, C.L., Abe, T., Tamaki, S., Lin, Y.W., et al. (2020). Behavioral and brain-transcriptomic synchronization between the two opponents of a fighting pair of the fish Betta splendens. PLoS Genet 16(6), e1008831. DOI:10.1371/journal.pgen.1008831.

Wang, Y., and Sun, Z. (2009). Current understanding of klotho. Ageing Res Rev 8(1), 43–51. DOI:10.1016/j.arr.2008.10.002.

Wang, Z., Qi, F., Cui, Y., Zhao, L., Sun, X., Tang, W., and Cai, P. (2018). An update on Chinese herbal medicines as adjuvant treatment of anticancer therapeutics. Biosci Trends 12(3), 220–239. DOI:10.5582/bst.2018.01144.

Weischenfeldt, J., Lykke-Andersen, J., and Porse, B. (2005). Messenger RNA surveillance: neutralizing natural nonsense. Curr Biol 15(14), R559–562. DOI:10.1016/j.cub.2005.07.002.

Wong, J.J., Ritchie, W., Ebner, O.A., Selbach, M., Wong, J.W., Huang, Y., Gao, D., Pinello, N., Gonzalez, M., Baidya, K., et al. (2013). Orchestrated intron retention regulates normal granulocyte differentiation. Cell 154(3), 583–595. DOI:10.1016/j.cell.2013.06.052.

Yabaluri, N., and Bashyam, M.D. (2010). Hormonal regulation of gluconeogenic gene transcription in the liver. J Biosci 35(3), 473–484. DOI:10.1007/s12038-010-0052-0.

Yeo, G., and Burge, C.B. (2004). Maximum entropy modeling of short sequence motifs with applications to RNA splicing signals. J Comput Biol 11(2-3), 377–394. DOI:10.1089/1066527041410418.

Yoshioka, H., Fukaya, S., Tominaga, S., Nagatsu, A., Miura, N., and Maeda, T. (2019). Protective effect of the Kampo formula “Juzen-taiho-to” on isoniazid- and rifampicin-induced hepatotoxicity in mice. Fundamental Toxicological Sciences 6(1), 25–29. DOI:10.2131/fts.6.25.

Zane, L., Sharma, V., and Misteli, T. (2014). Common features of chromatin in aging and cancer: cause or coincidence? Trends Cell Biol 24(11), 686–694. DOI:10.1016/j.tcb.2014.07.001.

Zelcer, N., and Tontonoz, P. (2006). Liver X receptors as integrators of metabolic and inflammatory signaling. J Clin Invest 116(3), 607–614. DOI:10.1172/JCI27883.

Zhou, X., Seto, S.W., Chang, D., Kiat, H., Razmovski-Naumovski, V., Chan, K., and Bensoussan, A. (2016). Synergistic Effects of Chinese Herbal Medicine: A Comprehensive Review of Methodology and Current Research. Front Pharmacol 7, 201. DOI:10.3389/fphar.2016.00201.

