## Supplementaly file for "Intron retention as a new pre-symptomatic marker of aging and its recovery to the normal state by a traditional Japanese multi-herbal medicine"

### Supplements

### Fig. S1

(A)

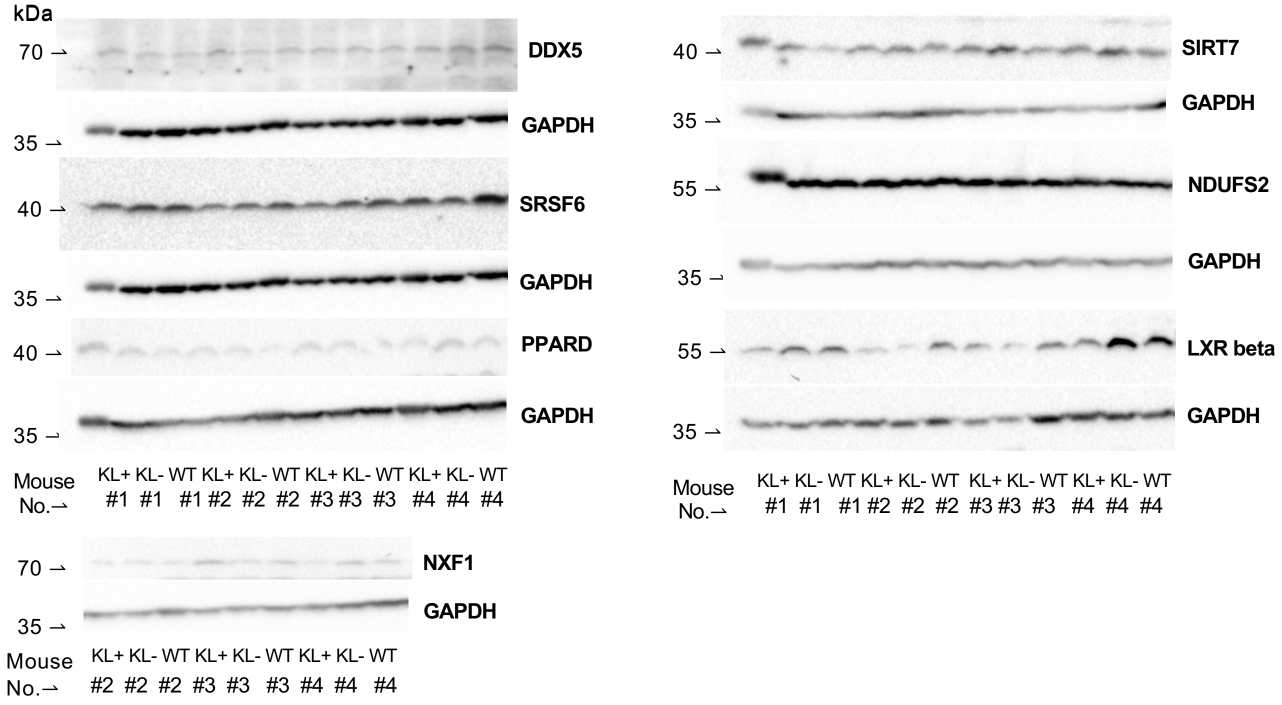


(B)

| | KL+ #1 | KL- #1 | WT #1 | KL+ #2 | KL- #2 | WT #2 | KL+ #3 | KL- #3 | WT #3 | KL+ #4 | KL- #4 | WT #4 |
|---|---|---|---|---|---|---|---|---|---|---|---|---|
| **DDX5** | 631384 | 390864 | 301403.7 | 660624.7 | 363636.5 | 610454.8 | 531780.3 | 470939.4 | 460357.1 | 399428 | 727991 | 828326.8 |
| **GAPDH** | 503238.9 | 946792 | 1146939 | 1008476 | 851865 | 998416 | 739674.9 | 597647.2 | 932221.6 | 891783.4 | 1147271 | 1002109 |
| **DDX5/GAPDH** | 1.254641 | 0.41283 | 0.26279 | 0.655073 | 0.426871 | 0.611423 | 0.718938 | 0.787989 | 0.493828 | 0.447898 | 0.634542 | 0.826584 |

| | KL+ #1 | KL- #1 | WT #1 | KL+ #2 | KL- #2 | WT #2 | KL+ #3 | KL- #3 | WT #3 | KL+ #4 | KL- #4 | WT #4 |
|---|---|---|---|---|---|---|---|---|---|---|---|---|
| **SRSF6** | 256135.3 | 304260.9 | 362489.7 | 69105.06 | 167561.3 | 264056 | 166504 | 247421.7 | 365182.9 | 372849.2 | 274956.3 | 1164866 |
| **GAPDH** | 503238.9 | 946792 | 1146939 | 1008476 | 851865 | 998416 | 739674.9 | 597647.2 | 932221.6 | 891783.4 | 1147271 | 1002109 |
| **SRSF6/GAPDH** | 0.508974 | 0.32136 | 0.31605 | 0.068524 | 0.196699 | 0.264475 | 0.225104 | 0.413993 | 0.391734 | 0.418094 | 0.239661 | 1.162415 |

| | KL+ #1 | KL- #1 | WT #1 | KL+ #2 | KL- #2 | WT #2 | KL+ #3 | KL- #3 | WT #3 | KL+ #4 | KL- #4 | WT #4 |
|---|---|---|---|---|---|---|---|---|---|---|---|---|
| **PPARD** | 15554.26 | 7908.874 | 4765.61 | 5580.388 | 5339.924 | 3780.64 | 6898.803 | 6623.459 | air | 6360.702 | 11479.67 | 6750.317 |
| **GAPDH** | 4756.024 | 5244.539 | 3795.882 | 3130.64 | 3329.882 | 5534.024 | 5903.539 | 5576.711 | 7910.66 | 6225.317 | 7708.66 | 8530.61 |
| **PPARD/GAPDH** | 3.270433 | 1.508021 | 1.255468 | 1.782507 | 1.603638 | 0.683163 | 1.168588 | 1.1877 | - | 1.021747 | 1.489192 | 0.791305 |

| | KL+ #1 | KL- #1 | WT #1 | KL+ #2 | KL- #2 | WT #2 | KL+ #3 | KL- #3 | WT #3 | KL+ #4 | KL- #4 | WT #4 |
|---|---|---|---|---|---|---|---|---|---|---|---|---|
| **SIRT7** | 9860.823 | 5874.439 | 3422.418 | 5942.489 | 7087.267 | 6450.51 | 7923.095 | 12057.05 | 5781.317 | 8270.631 | 11546.05 | 6843.024 |
| **GAPDH** | 4003.832 | 10502.1 | 4715.246 | 5377.368 | 9363.388 | 10548.92 | 5852.368 | 4523.539 | 4440.882 | 3244.296 | 3466.64 | 7273.196 |
| **SIRT7/GAPDH** | 2.462846 | 0.559359 | 0.72582 | 1.105092 | 0.756913 | 0.611485 | 1.353827 | 2.665401 | 1.30184 | 2.549284 | 3.330616 | 0.940855 |

| | KL+ #1 | KL- #1 | WT #1 | KL+ #2 | KL- #2 | WT #2 | KL+ #3 | KL- #3 | WT #3 | KL+ #4 | KL- #4 | WT #4 |
|---|---|---|---|---|---|---|---|---|---|---|---|---|
| **NDUFS2** | 3315.589 | 6451.64 | 3846.811 | 3678.518 | 4673.811 | 3863.225 | 4029.811 | 3508.811 | 3561.983 | 2587.397 | 2366.447 | 2387.447 |
| **GAPDH** | 7404.095 | 5752.853 | 7529.146 | 10785.34 | 12529.17 | 11255.87 | 11898.87 | 10320.46 | 10726.17 | 8631.167 | 9966.045 | 9466.167 |
| **NDUFS2/GAPDH** | 0.447805 | 1.121468 | 0.510923 | 0.341067 | 0.373034 | 0.343219 | 0.338672 | 0.339986 | 0.332083 | 0.299774 | 0.237451 | 0.252208 |

| | KL+ #1 | KL- #1 | WT #1 | KL+ #2 | KL- #2 | WT #2 | KL+ #3 | KL- #3 | WT #3 | KL+ #4 | KL- #4 | WT #4 |
|---|---|---|---|---|---|---|---|---|---|---|---|---|
| **LXR beta** | 1730.497 | 3649.175 | 3380.054 | 1586.134 | 891.92 | 2703.397 | 2428.69 | 1159.82 | 3600.225 | 4177.711 | 10825.44 | 8957.924 |
| **GAPDH** | 2685.933 | 3322.175 | 4227.004 | 5012.246 | 4353.054 | 4564.418 | 2740.225 | 2457.569 | 7461.418 | 7465.317 | 7297.953 | 5791.832 |
| **LXR beta/GAPDH** | 0.644282 | 1.098429 | 0.799633 | 0.316452 | 0.204895 | 0.592276 | 0.88631 | 0.471938 | 0.482512 | 0.559616 | 1.483353 | 1.546648 |

| | KL+ #2 | KL- #2 | WT #2 | KL+ #3 | KL- #3 | WT #3 | KL+ #4 | KL- #4 | WT #4 |
|---|---|---|---|---|---|---|---|---|---|
| **NXF1** | 2075.861 | 2581.518 | 2711.69 | 7328.004 | 3699.619 | 4504.589 | 2624.468 | 5605.711 | 3741.983 |
| **GAPDH** | 4886.125 | 4245.246 | 6572.439 | 4580.246 | 4770.368 | 5674.368 | 5823.368 | 7943.317 | 9783.782 |
| **NXF1/GAPDH** | 0.424848 | 0.608096 | 0.412585 | 1.599915 | 0.775542 | 0.793849 | 0.450679 | 0.705714 | 0.382468 |

(C)

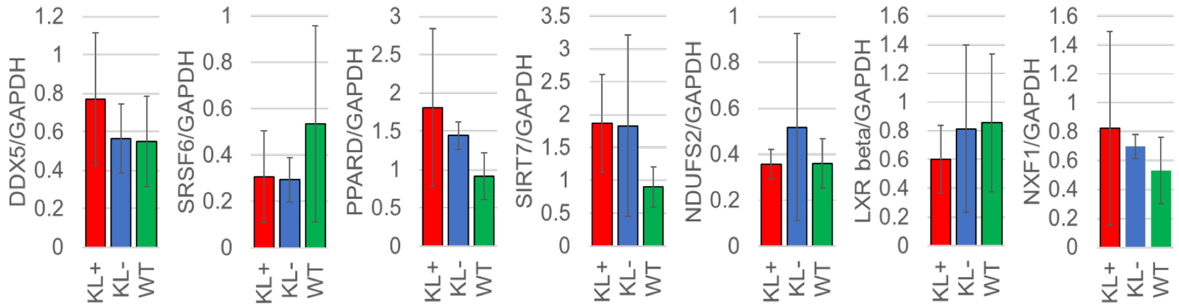


**Figure S1. Characterization of proteins.** (A) Images of seven representative membranes by using Western blotting were shown. GAPDH used as a control. (B) Tables showed intensity values of each band. They were normalized by using those of GAPDH. (C) Bar plots showed quantitation of protein revels. One-way ANOVA was used to calculate statistical significance. Expression levels did not change in KL+, KL–, and WT. Error bars indicate mean ± standard deviations (n =3 or 4).

**Fig. S2**

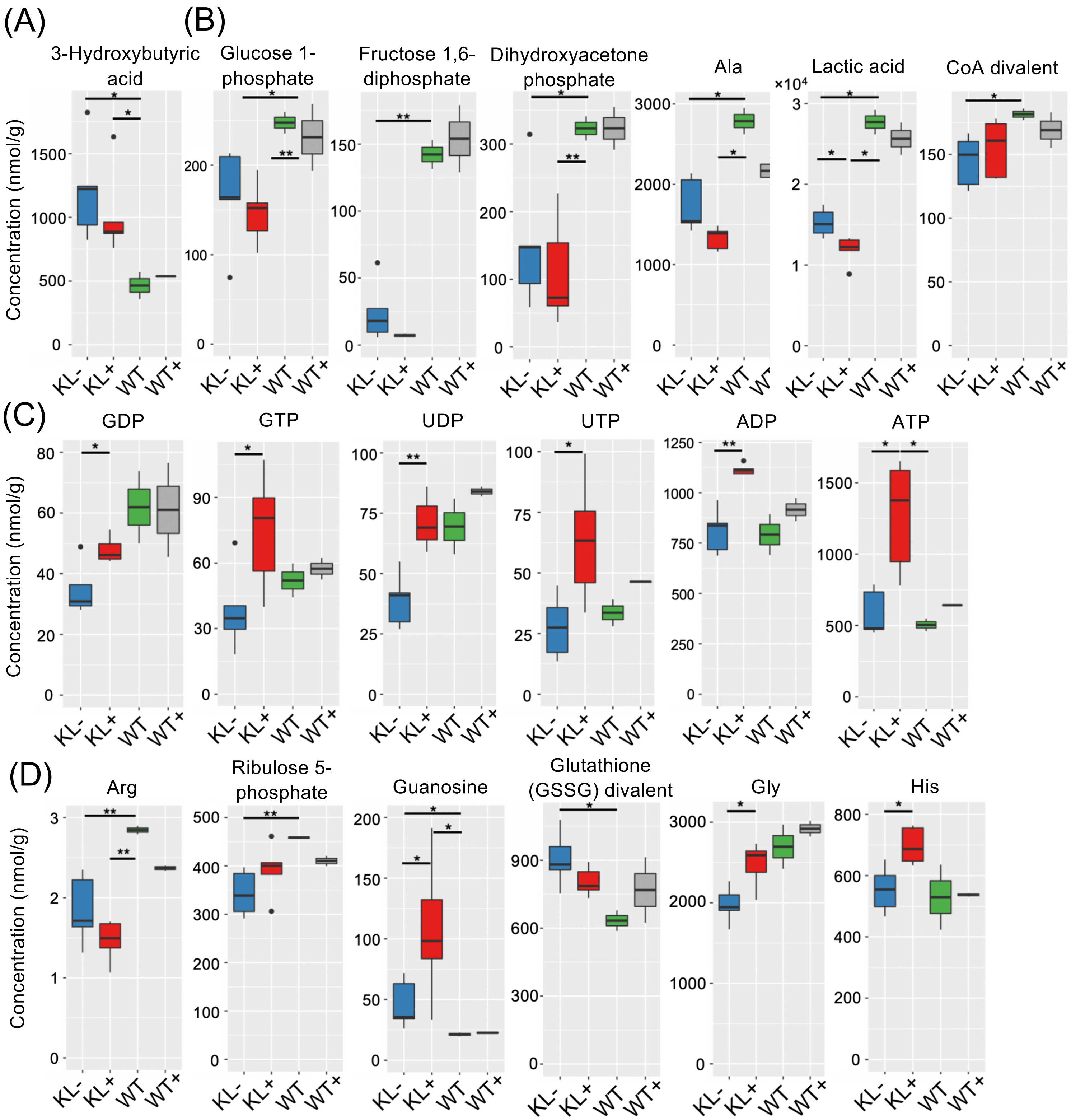


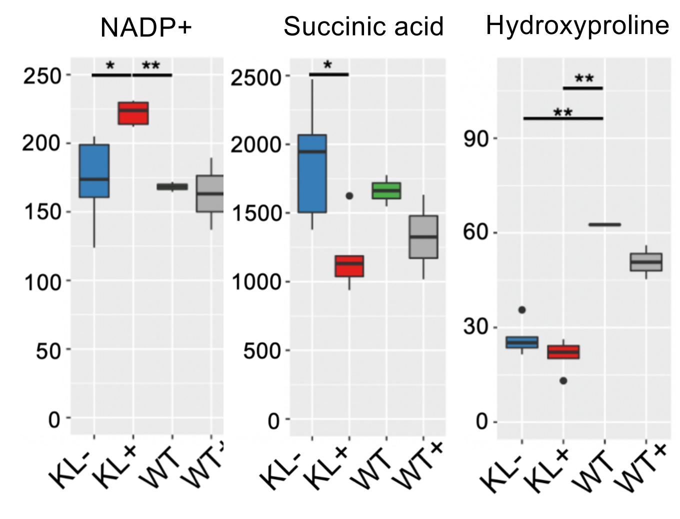


**Figure S2. A metabolome analysis showed that the klotho mice are undergoing a starvation-like condition at 7 weeks of age.** Based on a metabolome analysis of (A) 3-HBA and (B) other glycolysis-related metabolites, KL– data indicate that these mice are undergoing a starvation-like condition. (C) Based on the metabolome analysis about metabolites related to RNA precursors, the concentrations of some nucleotides were recovered in KL+ in comparison with KL– and WT. Data in (D) are those of 9 additional metabolites. **P* < 0.05, ***P* < 0.01, unpaired Student's *t*-test.

**Fig. S3**

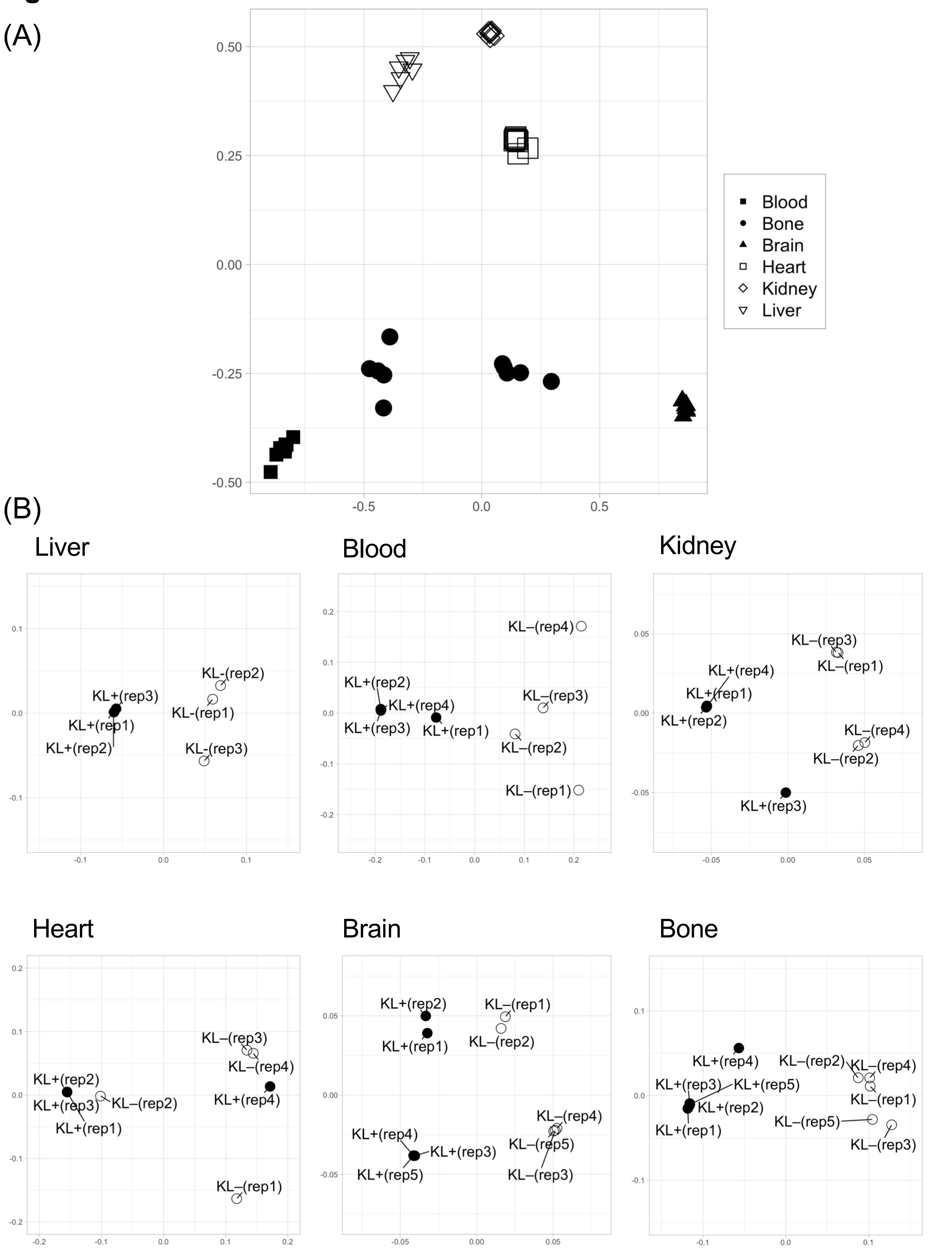


**Fig. S3. Multi-dimensional scaling of transcriptomes of 6 organs in klotho mice.** (A) shows well separation to one another, reflecting each organ's integrity in terms of RNA sequencing and those of (B) liver, blood, kidney, heart, brain and bone in klotho mice with (filled circles) or without (open circles) JTT.

### Fig.S4

(A)

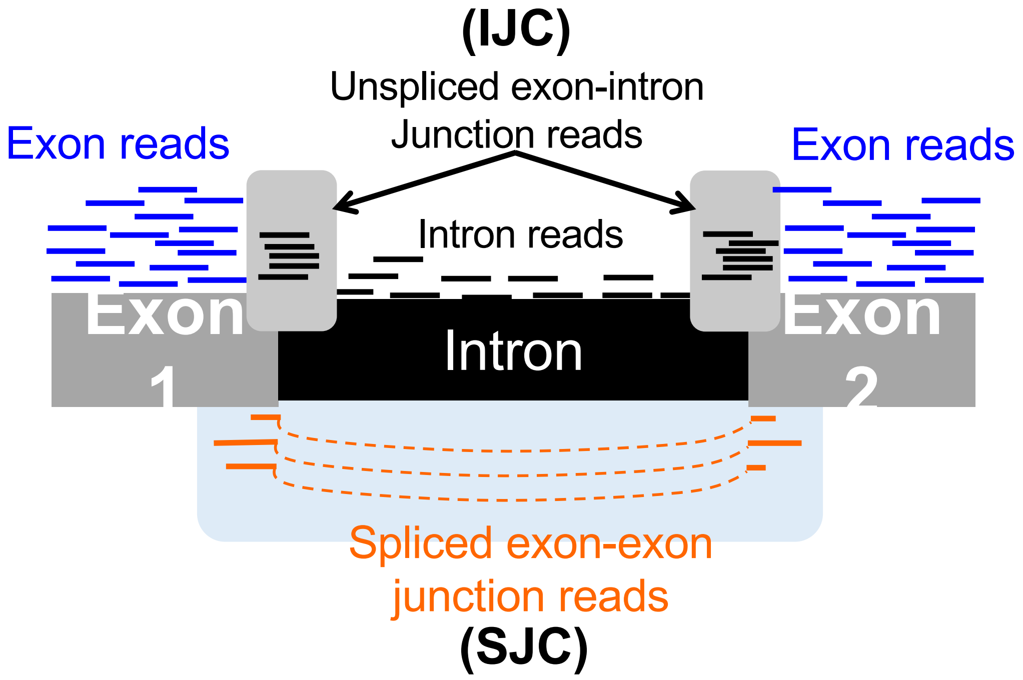


(B)

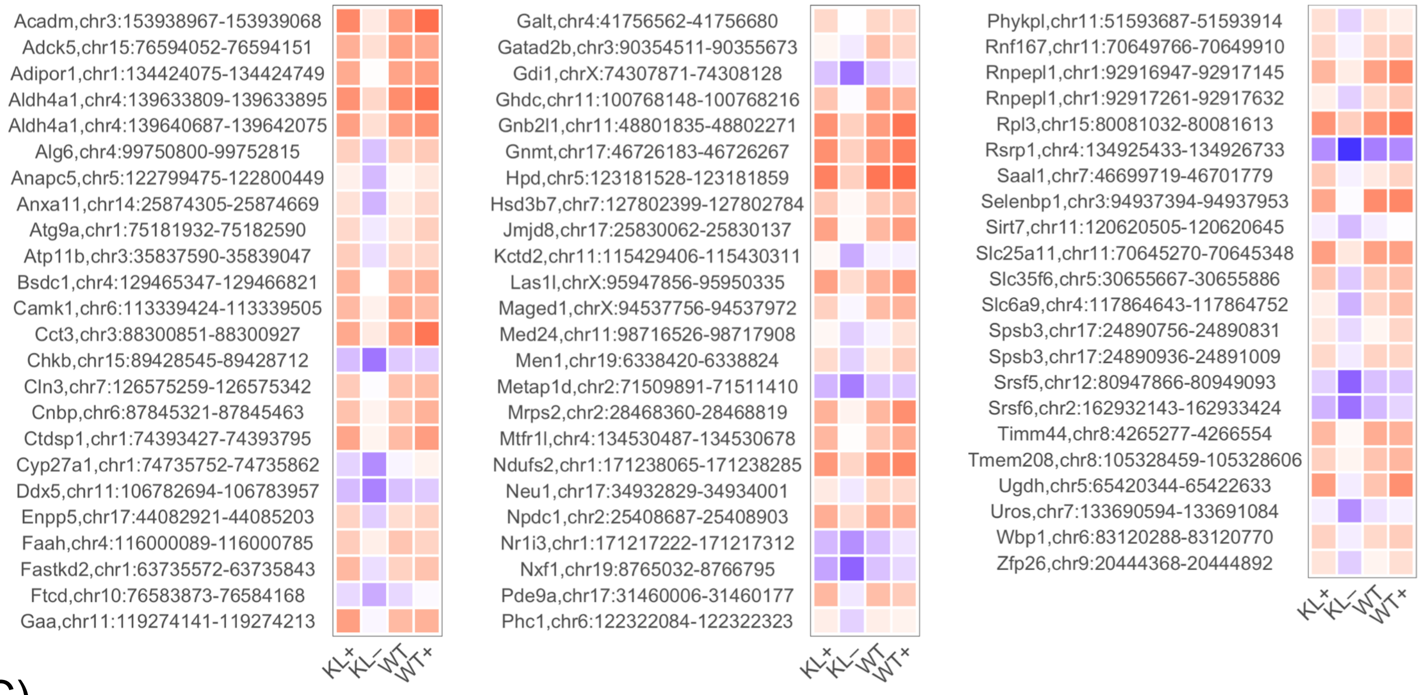


(C)

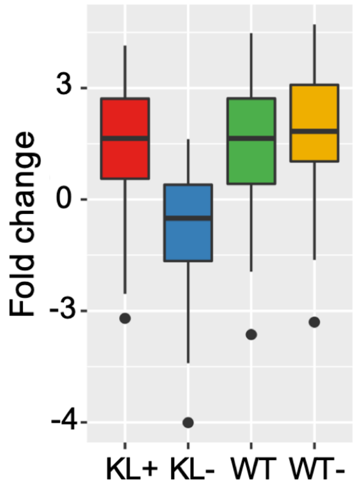


**Fig.S4. The extent of IR in WT with JTT is almost the same as that of WT in seventy loci of complete recovery.** (A) The quantification of IR events from mRNA-seq data using rMATS. The IJCs represent the reads containing the intron sequence at the junction. The SJCs represent the reads without intron sequences at the junction. (B) The data is the same as Fig.5BC except for the addition of WT+. Heat map illustration of FC values of loci in KL+, KL–, WT, and WT+ for the 70 “complete recovery” loci. (C) Boxplots of the FC values of the 70 “complete recovery” loci.

### Fig.S5

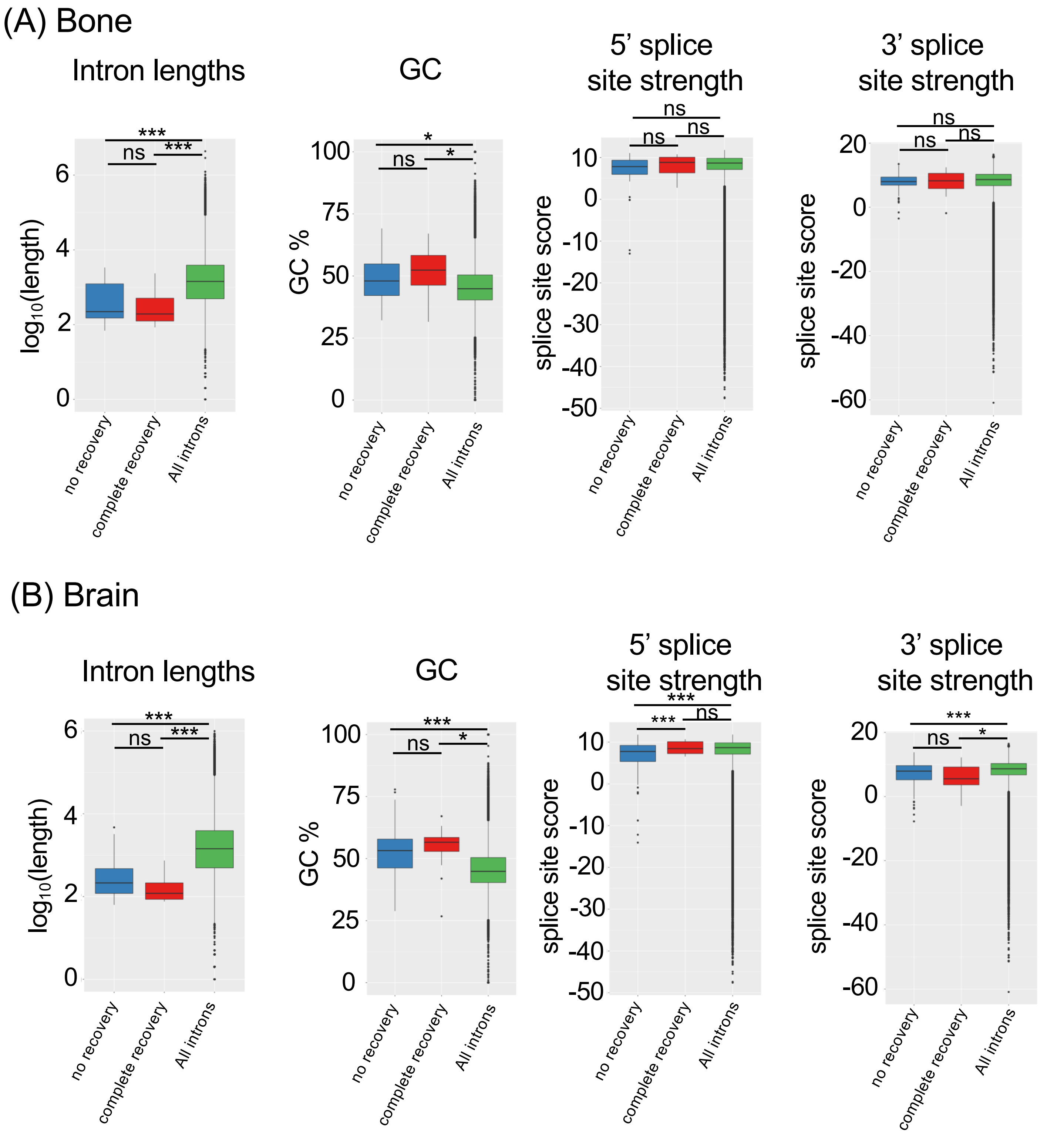


**Fig.S5 IR loci have their own characteristics.**
Boxplots showing intron lengths, GC % of intron sequences, and the strength score of 5'/3' splice sites compared among three groups of introns, namely "no recovery", "complete recovery" and "All introns" (254,005 loci) in bone (A) and brain (B). Statistical analyses are unpaired Student's t tests, and significance is annotated as $*P \leq 0.05$ , $**P \leq 0.01$, $***P \leq 0.001$.

**Fig. S6**

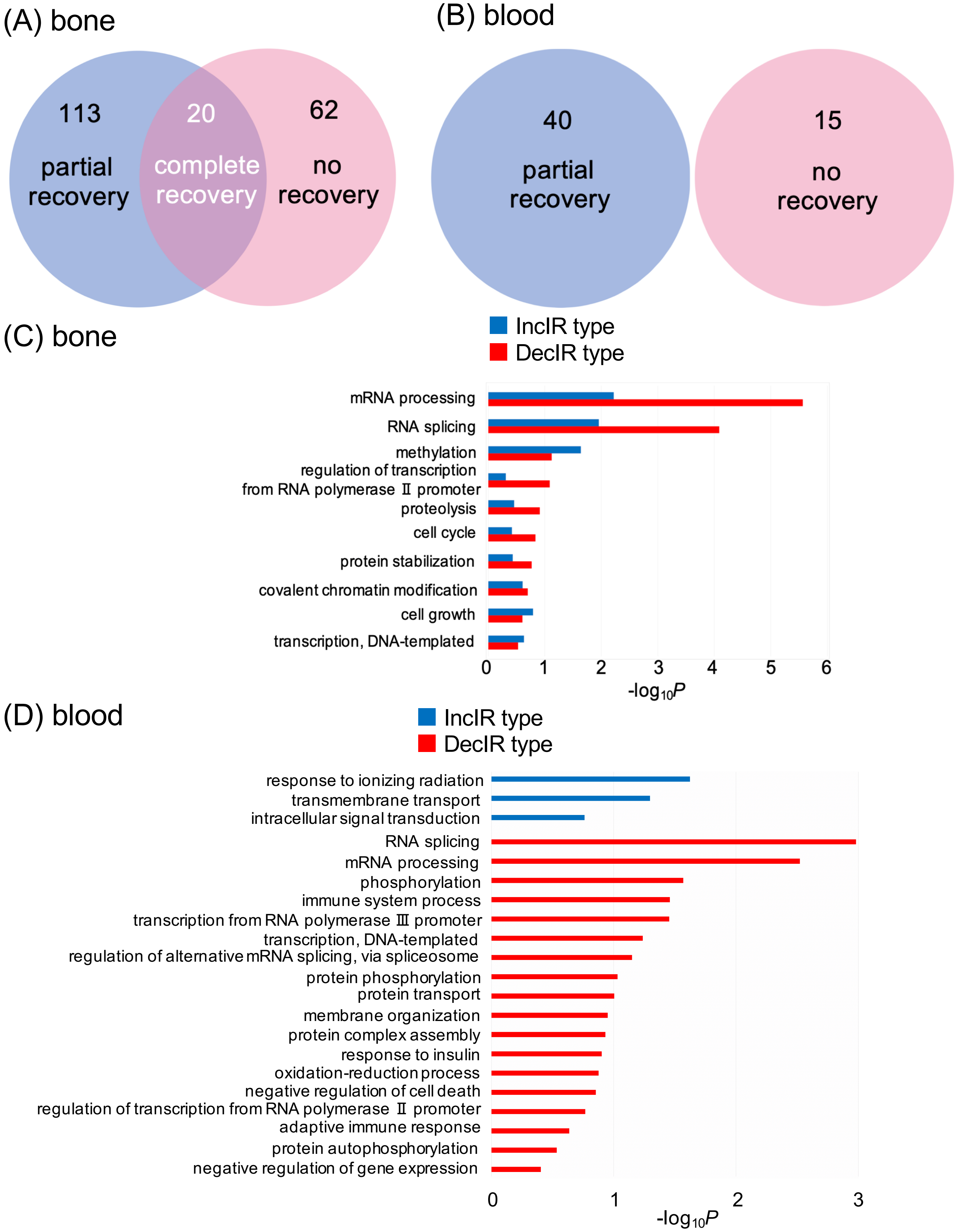


**Fig. S6 Quantification of DecIR and IncIR in bone and blood.** A Venn diagram of two types of loci, the DecIR type (blue circle) in the comparison of KL+ and KL– and the IncIR type (red circle) in the comparison of KL– and WT in bone (A) and blood (B). Bar graphs showing -$\log_{10}P$ of GO terms enriched for genes whose IR events were significantly changed in the bone (C) and blood (D).

**Fig.S7**

(A)

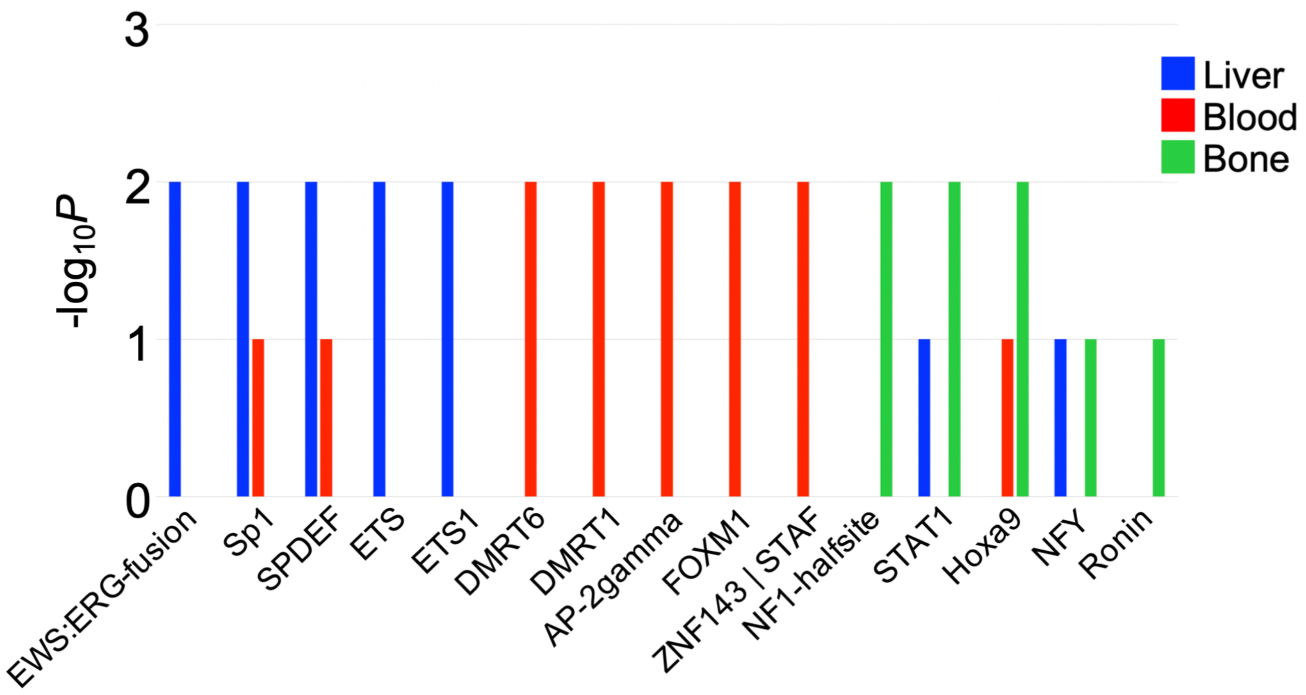


(B)

IncIR, Liver

175/179 genes, 190 loci

| TF | num of genes |
|---|---|
| KLF14 (Zf) | 120 |
| SCL (bHLH) | 93 |
| THRb (NR) | 93 |
| ETV4 (ETS) | 80 |
| Overall | 175 |

IncIR, Blood

13/14 genes, 15 loci

| TF | num of genes |
|---|---|
| KLF5 (Zf) | 9 |
| Sp1 (Zf) | 9 |
| HIF-1b (bHLH) | 6 |
| Overall | 13 |

IncIR, Bone

68/77 genes, 82 loci

| TF | num of genes |
|---|---|
| KLF14 (Zf) | 47 |
| SCL (bHLH) | 39 |
| Zac1 (Zf) | 33 |
| Overall | 68 |

**Fig S7. A very few TFs may control genes with IncIR.** (A) Bar graph showing TF-binding motifs that are significantly enriched in genes with IncIR from liver, blood and bone. (B) The table shows the combinations of TF- binding motifs that can cover more genes in the liver, blood and bone respectively.

**Table S1**. Primer sets of reverse transcription PCR

| gene | forward primer | reverse primer |
|---|---|---|
| *Acadm* | GTT CCC AGG CTC TTT TGA TG | GCA AGT TTG CCA GAG AGG AG |
| *Cdk11b* | GGG TGA TGA AAA GGA CTC TTG G | ACT TCA CCG AAG AAG CGT TG |
| *Cyp27a1* | TGC CTG AAA CCC TCC ATT CC | GCC GAT GGT CTC CTG AGT AC |
| *Ddx5* | GTG CCT GTT TTG GTA CTG CG | TAC CGA TGT GGC CTC CAG AG |
| *Decr2* | ATG TAG CTG TAG CAC CTG CC | AAC TTC CTAT GCC CTG CCA G |
| *Gnmt* | CTG AAG CCA GGA GAG CCA TC | ACC GGC TGG CAC TAA AGA AC |
| *Hpd* | TTT CCC TTG CTT GAT GAC GTG | CCT GAG AGA GGC CGG TTC C |
| *Hsd3b7* | TGG TCT ACA CGA GCA GCA TG | TTC CTT CCA TTG GCC TCG AG |
| *Itih3* | TCA ATA CCC TTC AGC AGC CC | CCT AGT TCA AGC CAC CCC TG |
| *Mug1* | GAT GAG TGT GCA GCT GGA AG | TCA GGA CTT TGA CTA CTG TGT CC |
| *Pcyt2* | ATG ACG TAG GGC CTC TTG G | CTG GTG CCT TTG ACCT GTT C |
| *Sirt7* | TGT AGA GTT TGG GTC GAC GG | GCA CCT CCT GCA TCC CTA AC |
| *Srsf5* | CCT CAA GAG TCA GCT GGC AG | TCC GCA AAG GTT ACT TCC CC |
